# HNRNPH1 regulates the neuroprotective cold-shock protein RBM3 expression through poison exon exclusion

**DOI:** 10.1101/2022.10.27.514062

**Authors:** Julie Qiaojin Lin, Deepak Khuperkar, Sofia Pavlou, Stanislaw Makarchuk, Nikolaos Patikas, Flora C.Y. Lee, Jianning Kang, Sarah F. Field, Julia M. Zbiegly, Joshua L. Freeman, Jernej Ule, Emmanouil Metzakopian, Marc-David Ruepp, Giovanna R. Mallucci

## Abstract

Enhanced expression of the cold-shock protein RNA binding motif 3 (RBM3) is highly neuroprotective both *in vitro* and *in vivo*. Whilst upstream signalling pathways leading to RBM3 expression have been described, the precise molecular mechanism of RBM3 induction during cooling remains elusive. To identify temperature-dependent modulators of RBM3, we performed a genome-wide CRISPR-Cas9 knockout screen using RBM3-reporter human iPSC-derived neurons. We found that RBM3 mRNA and protein levels are robustly regulated by several splicing factors, with heterogeneous nuclear ribonucleoprotein H1 (HNRNPH1) being the strongest positive regulator. Splicing analysis revealed that moderate hypothermia significantly represses the inclusion of a poison exon, which, when retained, targets the mRNA for nonsense-mediated decay. Importantly, we show that HNRNPH1 mediates this cold-dependent exon skipping via its interaction with a G-rich motif within the poison exon. Our study provides novel mechanistic insights into the regulation of RBM3 and provides further targets for neuroprotective therapeutic strategies.

**Graphical Abstract:** 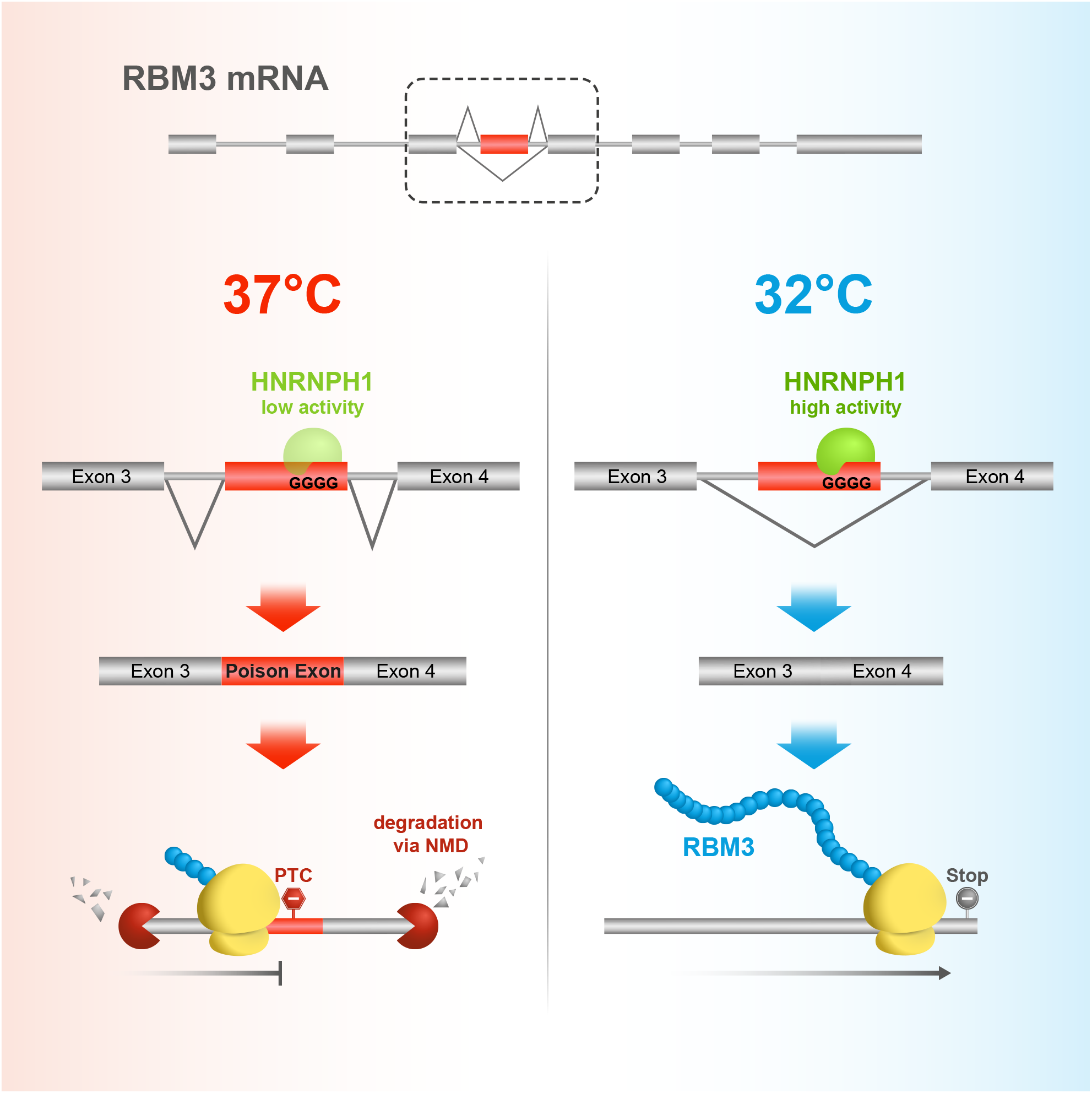

## Introduction

Lowering brain temperature during therapeutic hypothermia to ~35°C is widely used for neuroprotection in clinical practice, including in birth asphyxia (Cornette, 2012), acute ischemic stroke (Hemmen and Lyden, 2009), traumatic brain injury (Chen et al., 2019) and cardiac arrest (Presciutti and Perman, 2022). While the mechanisms of hypothermia-induced neuroprotection in humans are not fully understood, the cold-shock protein RNA-binding motif 3 (RBM3) is used as a biomarker of good outcome of therapeutic hypothermia in the intensive care setting (Ávila-Gómez et al., 2020; Rosenthal et al., 2019). In preclinical studies, RBM3 is neuroprotective in multiple settings. *In vivo,* increased RBM3 protein levels (induced by cooling or overexpression) restore memory, prevent synapse and neuronal loss and extend survival of prion and Alzheimer’s diseases (Bastide et al., 2017; Peretti et al., 2015, 2021); RBM3 stimulates neurogenesis in the rodent brain after hypoxic-ischemic brain injury and improves outcome (Zhu et al., 2019). *In vitro,* RBM3 protects neuronal cell lines from apoptosis during hypothermia (Chip et al., 2011). These findings have highlighted RBM3 as an attractive therapeutic target for neuroprotection - with the aim of inducing its expression without cooling. However, in this respect, a mechanistic understanding of its regulatory network is required.

In addition to hypothermia, RBM3 expression has been found to be modulated by hypoxia (Wellmann et al., 2004; Yan et al., 2019), serum starvation (Wellmann et al., 2010), metformin (Laustriat et al., 2015) and morphine exposure (Koo et al., 2012; Lefevre et al., 2020). Hypothermia induces RBM3 protein expression in association with TrkB activation *in vivo* (Peretti et al., 2021), but the exact molecular mechanism controlling RBM3 expression downstream of these signalling cascades and other modulators, remains unclear. Interestingly, hypothermia, metformin and hypoxia all alter genome-wide alternative splicing (Laustriat et al., 2015; Natua et al., 2021; Neumann et al., 2020) and differential splicing leading to altered mRNA and protein expression which is common among many RNA-binding proteins (Lareau et al., 2007; Müller-McNicoll et al., 2019; Sun et al., 2017). In particular, hypothermia-induced alternative splicing is observed in cold-inducible RNA-binding protein (CIRBP) transcripts (Gotic et al., 2016). These findings raise an interesting possibility that RBM3 gene expression could be fine-tuned on cooling in part by alternative splicing of its mRNA transcripts.

In this study, we uncovered the molecular mechanism involved in the cold induction of RBM3. An unbiased CRISPR/Cas9 whole-genome knockout screen in human induced pluripotent stem cell (iPSC)-derived neurons (i-neurons) identified several splicing factors, including heterogeneous nuclear ribonucleoprotein H1 (HNRNPH1), as trans-acting regulators of neuronal RBM3. We showed that HNRNPH1 mediates temperature-dependent RBM3 mRNA alternative splicing in multiple cell types, and maintains RBM3 transcript and protein expression in cooperation with the nonsense-mediated mRNA decay (NMD) pathway. Additionally, we located temperature-dependent cis-regulatory elements in the RBM3 mRNA and demonstrated that its functional interaction with HNRNPH1 is a key determinant of RBM3 differential splicing. These findings increase the range of therapeutic targets for RBM3 induction.

## Results

### Pooled CRISPR knockout screen identifies RNA splicing as a key regulatory pathway for RBM3

An RBM3 reporter iPSC line was generated by Cas9-mediated homology-directed repair to insert GFP at the N-terminus of the single copy of RBM3 on chromosome X in wild-type (WT) iPSCs, which contain a doxycycline (Dox)-inducible neurogenin 2 expression cassette (Pawlowski et al., 2017) and express Cas9 driven by the GAPDH promoter (Cas9 WT) (Fig. S1A). GFP-RBM3 predominantly localised to the nucleus in iPSCs (Fig. S1B) and i-neurons (Fig. 1A), consistent with previous reports (Rzechorzek et al., 2015). In this study, all Cas9-mediated gene editing in i-neurons was conducted by lentiviral transduction 4 days post differentiation, when Cas9 expression is optimal (Fig. S1A). Editing efficiencies were measured to be above 60% in GFP-RBM3 i-neurons 18 days post differentiation (Fig. S1C).

**Figure 1.**
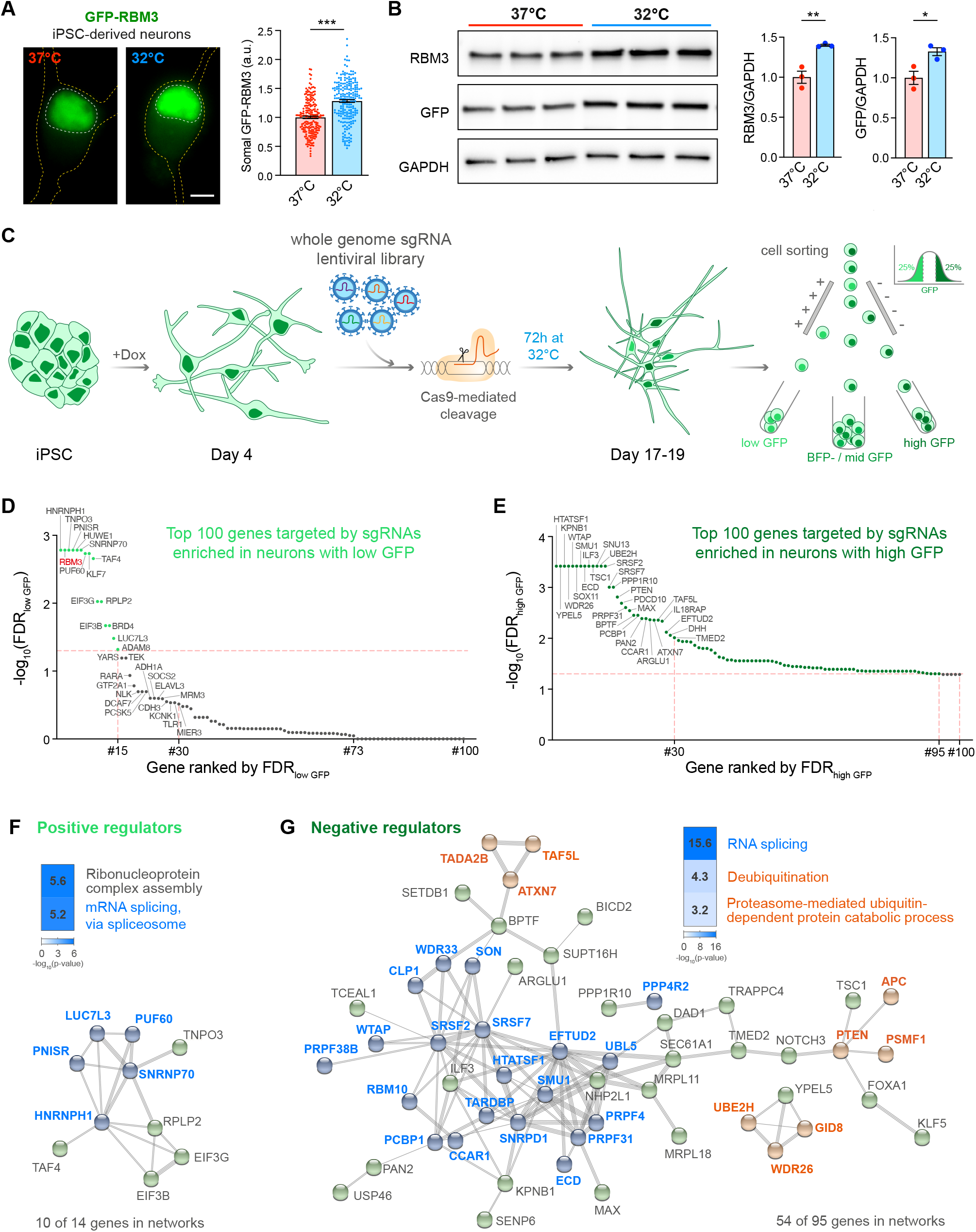
RBM3 CRISPR knockout screen identifies splicing factors as key RBM3 regulators. See also Fig. S1 and Table S2. (A) Representative images and quantification of somal intensity of GFP-RBM3 i-neurons at 37°C or after 72h cooling at 32°C. Nuclei and cells are outlined by white and yellow dashed lines, respectively. N= 3 (B) Western blots and quantification of RBM3 and GFP normalised to GAPDH in GFP-RBM3 i-neurons at 37°C or 32°C (72h). (C) Schematic of experimental steps in RBM3 CRISPR screen in i-neurons. GFP-RBM3 iPSCs stably expressing Cas9 after 4 days of Dox-induced differentiation are transduced with a whole-genome lentiviral sgRNA library expressing a BFP reporter. 10-12 days after transduction, the i-neuron cultures are incubated at 32°C for 72h, followed by FACS to sort BFP-positive i-neurons with the highest and lowest 25% GFP fluorescence intensity into separate pools. N = 2 GFP-RBM3 clones and 3 biological replicates. (D-E) Top 100 RBM3 positive regulator candidates with their sgRNAs enriched in the low-GFP i-neuron pool (D). Top 100 RBM3 negative regulator candidates with their sgRNAs enriched in the high-GFP i-neuron pool (E). Genes ranked by statistical significance (FDR). Horizontal dashed line: FDR=0.05. (F) The top-ranked Gene Ontology terms and STRING networks of 14 positive regulator candidates (FDR<0.05). Genes related to RNA splicing are indicated in blue. (G) The top-ranked Gene Ontology terms and STRING networks of 95 positive regulator candidates (FDR<0.05). Genes related to RNA splicing are coloured in blue. Genes involved in deubiquitination or proteasome-mediated ubiquitin-dependent protein catabolic processes are coloured in orange. Mean ± SEM; * (p<0.5), **(p < 0.01), ***(p < 0.001); unpaired t-tests. Scale bars: 5 μm.

GFP fluorescence intensities in both clones of GFP-RBM3 i-neurons were significantly enhanced by cooling at 32°C for 72h (Fig. 1A, S1D) as a result of increased nuclear and cytosolic GFP-RBM3 levels (Fig. S1E). Around 30% cold-induction of GFP-RBM3 protein levels was also shown by western blots (Fig. 1B). These observations validated that endogenously GFP-tagged RBM3 is temperature-sensitive, consistent with behaviours of unmodified RBM3 both seen with the Cas9 WT cells in this study and reported RBM3 hypothermic induction (Jackson et al., 2015; Peretti et al., 2015).

Using both clones of this characterised GFP-RBM3 reporter line, we performed an RBM3 CRISPR knockout pooled screen (Fig. 1C) by transducing 4-day differentiated cells with a custom-made whole-genome sgRNA lentiviral library, targeting critical exons of 18466 genes across the human genome, co-expressing a BFP reporter. Day 18 i-neurons were incubated at 32°C for 72h before dissociation and fluorescence-activated cell sorting (FACS). BFP-positive cells with the highest and lowest GFP-RBM3 expression, which fell in the top and bottom quartile of GFP intensity profiles, were separately collected, denoted as high GFP and low GFP populations. Genomic DNA was sequenced to identify sgRNAs enriched in high or low GFP populations (see Table S2). Ranked by the false discovery rate (FDR), 15 genes, including RBM3, were enriched in low GFP i-neurons, suggesting they likely positively regulate RBM3 expression (Fig. 1D). In contrast, 95 genes, likely to negatively affect RBM3 expression, were enriched in high GFP populations (Fig. 1E).

Gene ontology and network analysis revealed that a major group of RBM3 regulator candidates are involved in RNA splicing (Fig. 1F, G), including spliceosomal proteins, e.g., U1 small nuclear ribonucleoprotein 70 kDa (SNRNP70), heterogeneous nuclear ribonucleoproteins (hnRNPs) and serine and arginine rich splicing factors (SRSFs). Other potential RBM3 regulators play roles in nucleocytoplasmic transport, e.g. transportin 3 (TNPO3) and karyopherin-β1 (KPBN1); translation, e.g. 60S acidic ribosomal protein P2 (RPLP2) and eukaryotic translation initiation factor 3 subunit B (EIF3B); transcription, e.g. transcription initiation factor TFIID subunit 4 (TAF4); ubiquitination, e.g. ubiquitin-conjugating enzyme E2 H (UBE2H).

### Arrayed target validation reveals regulators modulating neuronal RBM3 protein and transcript abundance

To validate the top hits identified by the pooled screen, we individually depleted the top 30 RBM3 positive and negative regulator candidates using 1-3 sgRNAs per gene from the whole-genome library (see Table S3). I-neurons transduced with 3 non-targeting sgRNAs from the pooled library or the reporter lentivirus for editing efficiency calculation served as controls. As a positive control, RBM3 sgRNAs reduced GFP fluorescence by over 80% compared to non-targeting sgRNAs at 37°C or 32°C (Fig. 2A). Moderate spectral crossover and activation of specific signalling pathways due to target-specific genome editing may account for the discrepancy between non-targeting sgRNA and reporter controls. To apply a more stringent standard, we performed statistical analysis between specific gene knockout (KO) groups and one of the two control groups with high p values, for instance, comparing to the non-targeting sgRNA control showing lower GFP intensity to identify positive regulators, and to the reporter control for negative regulators. Knocking out 7 out of 29 positive regulator candidates tested significantly reduced GFP-RBM3 levels at 37°C and 32°C (Fig. 2A). HNRNPH1 (Chou et al., 1999), TNPO3 (Kataoka et al., 1999), PNN Interacting Serine And Arginine Rich Protein (PNISR) (Zimowska et al., 2003) and SNRNP70 (Pomeranz Krummel et al., 2009) are associated with RNA splicing. Among 30 tested RBM3 negative regulator candidates, KO of 6 genes significantly increased GFP-RBM3 expression in i-neurons at 37°C and 32°C (Fig. 2B). Specifically, HIV-1 Tat Specific Factor 1 (HTATSF1) and WT1 Associated Protein (WTAP) are splicing regulators, while Yippee Like 5 (YEPL5), WD Repeat Domain 26 (WDR26) and UBE2H mediate ubiquitination and proteasomal degradation, the depletion of which is more likely to result in generic instead of RBM3-specific protein accumulation. Changes in GFP-RBM3 levels upon selective RBM3 regulator KO were also confirmed using fluorescent microscopy (Fig. S2A). Collectively, the arrayed CRISPR knockout assay validated hits associated with the lowest FDR in the pooled screen (Fig. 1D, E), and reiterated that multiple components of the splicing machinery are key to RBM3 protein expression.

**Figure 2.**
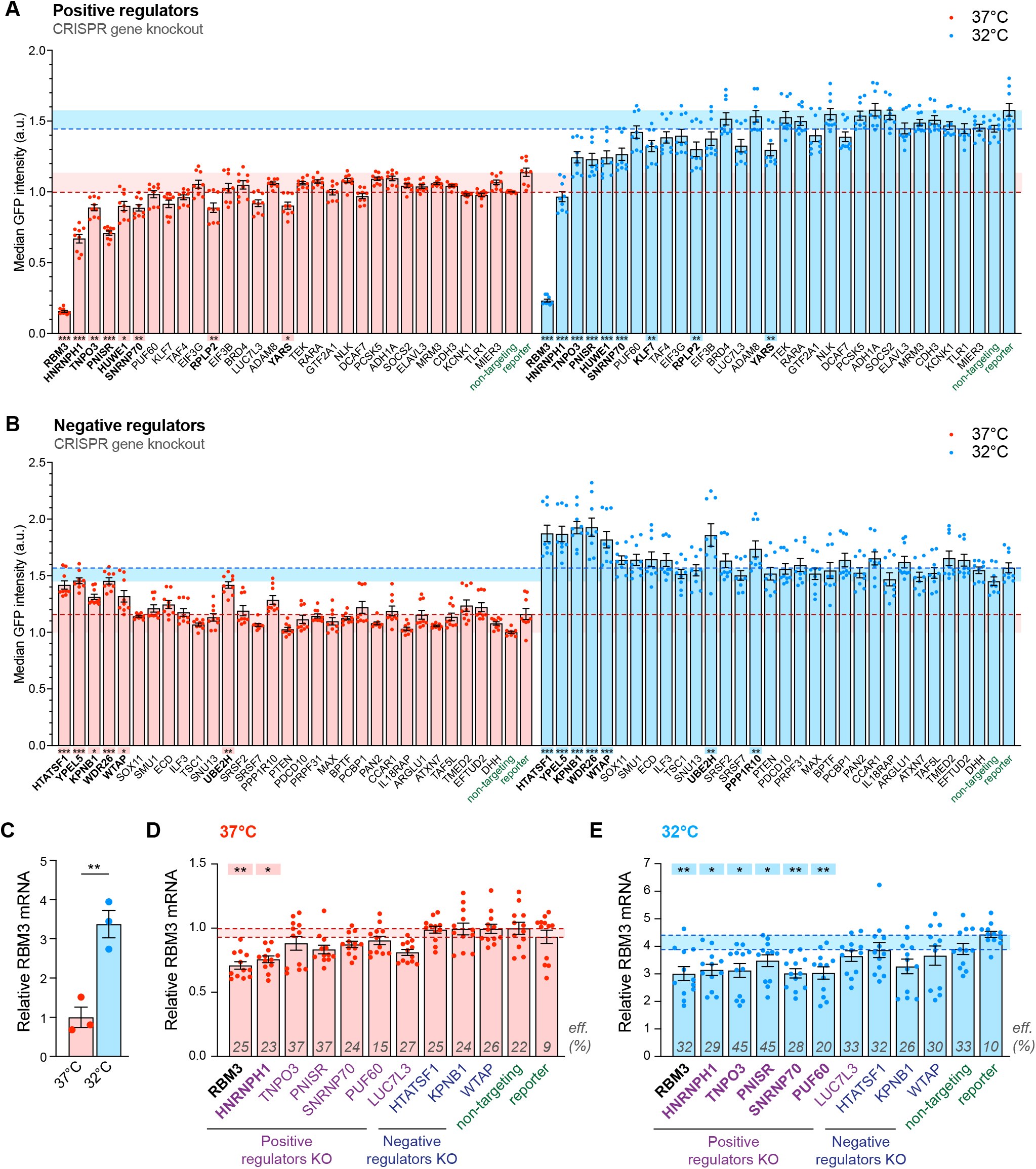
Depleting RBM3 positive regulators function in RNA splicing reduces RBM3 protein and mRNA levels. See also Fig. S2 and Table S3. (A-B) Median GFP intensity of BFP-positive GFP-RBM3 i-neurons measured by flow cytometry upon the sgRNA/Cas9-mediated KO of top 30 positive (A) or negative (B) regulator candidates. Statistical analysis is performed between the specific and non-targeting sgRNA groups for positive regulators (A) or between the specific sgRNA and reporter groups for negative regulators (B) within the 37°C or 32°C (72h) population. (C) qRT-PCR of RBM3 mRNA level normalised to 18s rRNA in i-neurons at 37°C or 32°C (72h). (D-E) qRT-PCR of RBM3 mRNA level normalised to 18s rRNA in GFP-RBM3 i-neurons at 37°C (D) or 32°C for 72h (E) transduced with lentivirus containing specific, non-targeting sgRNA or the reporter. Statistical analysis is performed between the specific and non-targeting sgRNA for negative regulators in (D) and positive regulators in (E), or between the specific and reporter groups for positive regulators in (D) and negative regulators in (E). Transduction efficiencies are indicated in corresponding bars. N≥3. Mean ± SEM; n.s. (not significant), * (p<0.5), **(p < 0.01), ***(p < 0.001); One-way ANOVA with multiple comparisons in (A), (B), (D), (E), unpaired t-tests in (C).

To investigate whether the cold-induced GFP-RBM3 protein expression is controlled at the translational or transcriptional level, the translational inhibitor cycloheximide or the RNA polymerase inhibitor actinomycin D was applied to i-neurons at 37°C or 32°C. The elevation of GFP-RBM3 in cooled cells was completely abolished by either inhibitor (Fig. S2B), indicating the cold-increased GFP-RBM3 expression relies on the *de novo* transcription and translation of RBM3 transcripts. In line with this finding, differential expression analysis comparing RNA-Seq data between i-neurons at 37°C and 32°C showed RBM3 increased 4-fold on cooling: this was the most significant increase across the entire transcriptome (Fig. S2C). Further, real-time PCR (quantitative PCR, qPCR) of RBM3 mRNA supports cold-induction of RBM3 mRNA by 3-5 folds in Cas9 WT i-neurons (Fig. 2C, S2D) and HeLa cells (Fig. S2E).

We next explored whether the changes of RBM3 protein expression upon individual regulator KO were due to altered RBM3 mRNA levels. We focused on the validated splicing regulating genes (HNRNPH1, PNISR, SNRNP70, HTATSF1, WTAP), potential regulators marginally affecting RBM3 expression (Poly(U) Binding Splicing Factor 60 (PUF60) and LUC7 Like 3 Pre-mRNA Splicing Factor (LUC7L3)) and the two nuclear protein importers (TNPO3 and KPNB1) (Fig. 2D, E). Due to cell death and promoter silencing, only 15-45% of the transduced cells remained BFP-positive, as approximates to transduction efficiencies (labelled on bars, Fig. 2D, E), on day 18 when RNA was extracted from i-neuron cultures at 37°C or 32°C. Remarkably, the extent of RBM3 transcript reduction due to HNRNPH1 KO at 37°C or 32°C was similar to RBM3 KO as a positive control (Fig. 2D, E), revealing HNRNPH1 as a strong positive regulator of RBM3 mRNA expression. TNPO3, PNISR, SNRNP70 and PUF60 KO also significantly lowered RBM3 transcript levels at 32°C (Fig. 2E), suggesting they are key regulators for cold induction of RBM3 transcripts. On the contrary, none of the tested negative regulator KO affected RBM3 mRNA expression, suggesting that their regulation may be at the translational or post-translational level. In support of our findings in i-neurons, RNA-Seq analysis of public data also showed reduced RBM3 mRNA levels upon HNRNPH1, SNRNP70 and PUF60 KD in K562 or HepG2 cells, while no difference was seen upon KPNB1 KD (Fig. S2F) (ENCODE Project Consortium, 2012; Luo et al., 2020). Particularly, both the mRNA and protein levels of RBM3 were most significantly reduced across the whole genome upon HNRNPH1 KD in HeLa cells (Uren et al., 2016), demonstrating the strong regulatory effect of HNRNPH1 on RBM3 transcripts and proteins. We further showed that HNRNPH1 KD significantly decreased RBM3 mRNA in HeLa cells by over 50% both at 37°C and 32°C (Fig. S2G). To summarise, depleting specific splicing factors, particularly HNRNPH1, abolishes the cold-induced increase of RBM3 mRNA expression in i-neurons and other cell types.

### RBM3 poison exon that triggers nonsense-mediated decay is silenced during hypothermia

Given that RBM3 mRNA expression is tightly regulated by splicing factors in i-neurons, we searched for RBM3 alternatively spliced isoforms with RNA-Seq data acquired from Cas9 WT i-neurons at 37°C and 32°C (Fig. 3A). Multiple isoforms of RBM3 transcripts were identified and the most abundant four isoforms could be distinguished by either Exon 3, Exon 3a-L or Exon 3a-S inclusion (Fig. 3A). Differential splicing analysis showed RBM3 Exon 3a inclusion levels drastically reduced upon cooling, from the percent spliced in (PSI) index of 0.023 to 0.002 for Exon 3a-L, or from 0.015 to 0.001 for Exon 3a-S (Fig. 3B, Table S4), while Exon 3 inclusion levels remained similar at 37°C and 32°C (Fig. S3A), demonstrating that cooling effectively repressed the production of Exon 3a-L/S-containing RBM3 transcripts.

**Figure 3.**
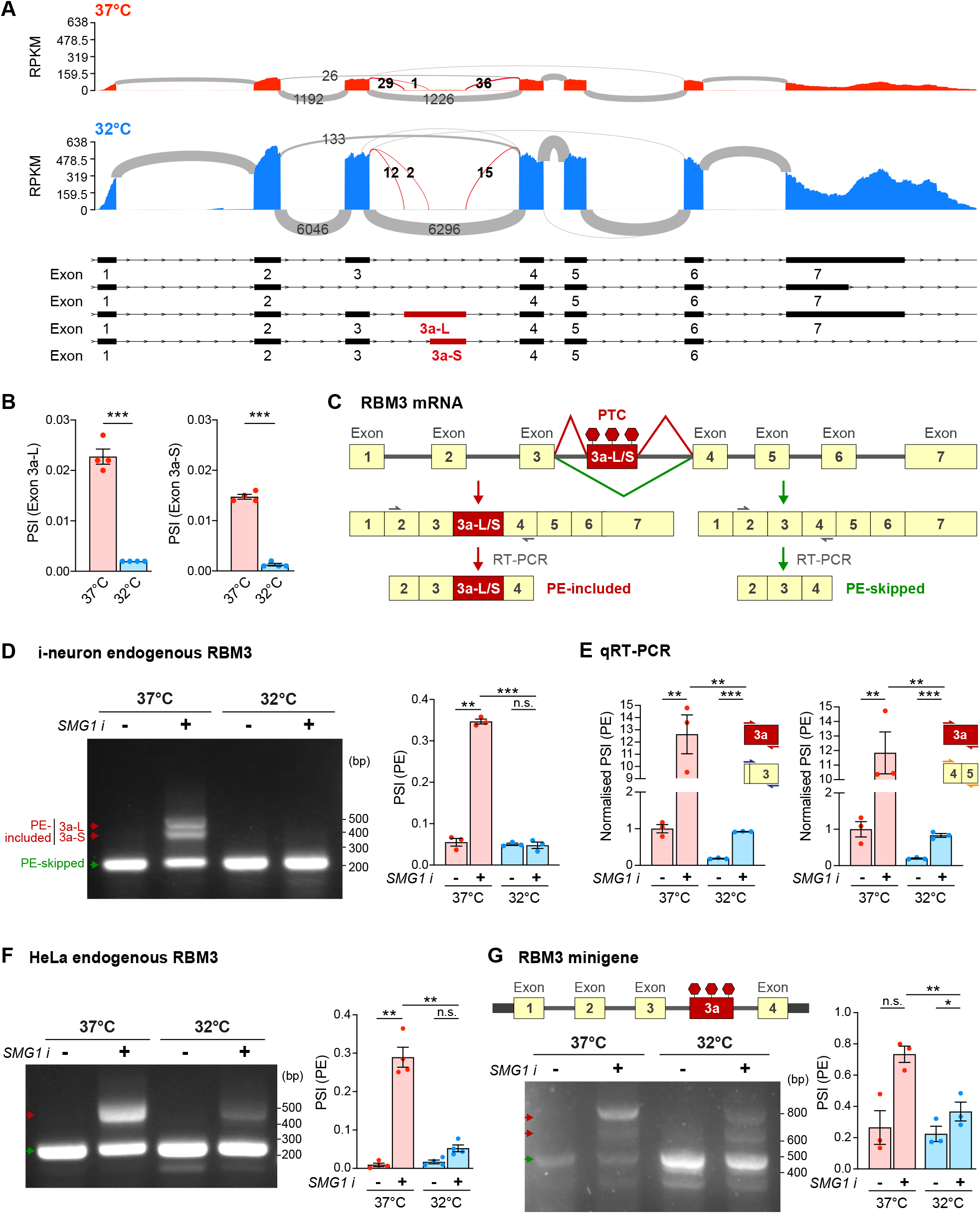
Hypothermia represses RBM3 mRNA poison exon inclusion. See also Fig. S3. (A) Sashimi plots of RBM3 transcripts in WT i-neurons at 37°C and 32°C (72h), showing major alternatively spliced isoforms. Differentially spliced Exon 3a-L and 3a-S junctions between 37°C and 32°C conditions are shown in red. N= 4. (B) PSI values of RBM3 Exon 3a-L and 3a-S relative to Exon 3 and 4 in i-neurons at 37°C or 32°C (72h). (C) Schematics of RBM3 Exon 3a, or poison exon (PE), alternative splicing and the resulting PE-included (left) or PE-skipped (right) mRNA products. RT-PCR primer pair amplifying Exon 2-4 are indicated by blue arrows. (D) RT-PCR of RBM3 mRNA (Exon 2-4) in i-neurons at 37°C or 32°C (72h) in the presence or absence of SMG1 inhibitor. PSI values of RBM3 PE are calculated based on the intensity of PE-included (red arrows) and PE-skipped (green arrow) isoforms. (E) qRT-PCR using a combination of primers targeting Exon 3a, Exon 3 or Exon 4-5 quantifies the PSI values of RBM3 PE (Exon 3a, including both 3a-L and 3a-S) relative to Exon 3 or Exon 4-5 at 37°C or 32°C (72h) in the presence or absence of SMG1 inhibitor. (F) RT-PCR of RBM3 mRNA (Exon 2-4) in HeLa cells at 37°C and 32°C (48h) in the presence or absence of SMG1 inhibitor. PSI values of RBM3 PE are depicted in the graph on the right. (G) Schematics and RT-PCR of RBM3 minigene (Exon 1-4), flanked by unique sequences (thick black bars) to distinguish it from endogenous transcripts during PCR amplification, expressed in HeLa cells at 37°C or 32°C (48h), in the presence or absence of SMG1 inhibitor. PSI values of RBM3 PE are depicted in the graph on the right. N≥3 Mean ± SEM; * (p<0.5), **(p < 0.01), ***(p < 0.001); FDR calculated by rMATS program in (B); unpaired t-tests in (E); paired t-test in (D), (F), (G).

A close examination of Exon 3a of RBM3 revealed multiple stop codons in-frame with the coding sequence, potentially leading to premature translational termination (Llorian et al., 2016). Thus, the inclusion of Exon 3a makes the RBM3 transcript a susceptible target for degradation by nonsense-mediated mRNA decay (NMD), an mRNA surveillance pathway that degrades mRNAs that harbour premature termination codons (PTCs) (He and Jacobson, 2015). Exon 3a could therefore serve as a poison exon (PE) in RBM3 mRNA, leading to a reduction of the transcript level if retained. Such post-transcriptional regulation mediated by PEs is key to fine-tuning expression levels for many proteins, especially RNA-binding proteins (Neumann et al., 2020). To further verify if Exon 3a indeed acts as a PE, and to explore the temperature-dependent inclusion of RBM3 PE quantitatively, we designed isoform-sensitive primers to amplify regions between RBM3 exon 3 and exon 4 to identify PE-included or PE-skipped isoforms of RBM3 transcripts (Fig. 3C). In order to detect the NMD-sensitive PE-included isoforms, we blocked NMD using a chemical inhibitor of SMG1 kinase, a key member of the NMD pathway (Langer et al., 2021). When i-neurons were incubated with SMG1 inhibitor for 24h at 37°C, both PE-included and PE-skipped isoforms were detected with RT-PCR. Interestingly, in 72h-cooled i-neurons, only the PE-skipped isoform was detected, even after SMG1 inhibitor treatment, clearly supporting that the RBM3 PE was preferentially excluded in cooled i-neurons (Fig. 3D). This finding was further validated by qPCR using primers against Exon 3a (detecting both 3a-L and 3a-S) and the constitutive exons, e.g., Exon 3 and Exon 4-5 (Fig. 3E). The fraction of PE-contained transcripts to the total transcripts at 37°C was over 5 times more compared to the fraction at 32°C. When NMD was inhibited by the SMG1 inhibitor, the differences rose to 13-15 folds (Fig. 3E).

Additionally, we designed an RBM3 minigene spanning RBM3 Exons 1-4, which was sensitive to NMD when expressed in HeLa cells (Fig. 3F). Consistent with the endogenous RBM3 mRNA (Fig. 3F), the minigene also showed a significantly lower fraction of PE-retained isoform at 32°C compared to 37°C, when NMD was blocked by SMG1 inhibitor (Fig. 3G) or cycloheximide (Fig. S3B).

### HNRNPH1 represses RBM3 poison exon inclusion at low temperatures

We next investigated whether positive RBM3 mRNA regulators control RBM3 transcript abundance through PE skipping. When NMD was inhibited, knocking down HNRNPH1 resulted in RBM3 PE retention in i-neurons, with 29% increase at 37°C and 57% increase at 32°C compared to the respective non-targeting control (Fig. 4A, B). RBM3 PE-retention upon HNRNPH1 KD was also found in published RNA-Seq analysis using K562 and HepG2 cells, demonstrating its conserved role across multiple cell types (Fig. 4C, S4A, S4B, Table S4) (ENCODE Project Consortium, 2012; Luo et al., 2020). In HeLa cells treated with SMG1 inhibitor, RNAi-mediated HNRNPH1 knockdown (Fig. S4C) increased PE retention in endogenous RBM3 transcripts (Fig. 4D) and in RBM3 minigene (Fig. 4E) only at 32°C when NMD was inhibited, suggesting that the role of HNRNPH1 in repressing RBM3 PE inclusion is context-dependent, being more important under the cooled condition. Moreover, overexpressing FLAG-tagged HNRNPH1 in HEK293T cells (Fig. S4D) repressed PE inclusion in endogenous RBM3 mRNA (Fig. 4F) and in the minigene (Fig. 4G) at 37°C, confirming the splicing-regulatory function of HNRNPH1 in RBM3 PE exclusion.

**Figure 4.**
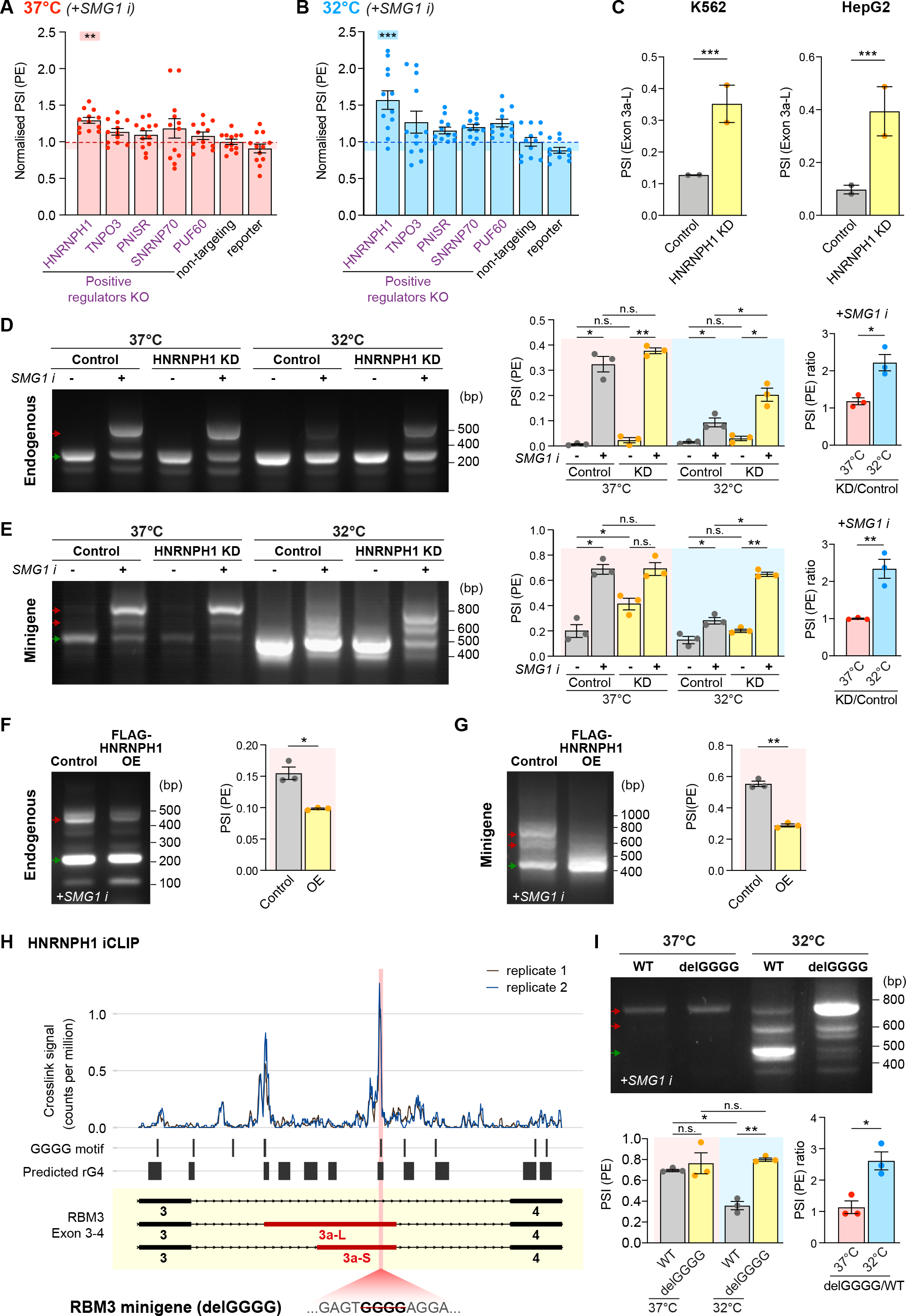
HNRNPH1 promotes RBM3 mRNA poison exon skipping. See also Fig. S4. (A-B) qRT-PCR quantifying the PSI values of RBM3 PE relative to RBM3 mRNA in WT i-neurons at 37°C (A) or 32°C (72h) (B), when NMD is blocked by SMG1 inhibitor. Statistical analysis is performed between the specific and non-targeting sgRNA groups. (C) PSI values of RBM3 Exon 3a-L in control and HNRNPH1-knocked down K562 and HepG2 cells. RNA-Seq data from ENCODE Project, 2 isogenic replicates are included. (D-E) RT-PCR of endogenous RBM3 (D) and expressed RBM3 minigene (E) in control (scramble siRNA) or HNRNPH1-knocked down HeLa cells at 37°C or 32°C (48h), in the presence or absence of SMG1 inhibitor. The ratio of PSI values between HNRNPH1 KD and control shown for only the SMG1 i-treated conditions. (F-G) RT-PCR of endogenous RBM3 (F) and expressed RBM3 minigene (G) in SMG1 inhibitor-treated control (untransfected) or FLAG-HNRNPH1-overexpressed (OE) HEK293T cells at 37°C. PSI values of RBM3 PE are shown in the graphs on the right respectively. (H) Analysis of public HNRNPH1 iCLIP dataset in two replicates, mapped to RBM3 Exon 3-4. Crosslink counts are normalised to library size. RNA G quadruplexes (rG4) are predicted using QGRS mapper. The position of the GGGG motif deleted in the mutant RBM3 minigene is shown in pink. (I) RT-PCR of WT and delGGGG RBM3 minigenes in HeLa cells at 37°C or 32°C (48h) treated with SMG1 inhibitor. PSI values of RBM3 PE are depicted in the adjacent graph N≥3 Mean ± SEM; * (p<0.5), **(p < 0.01), ***(p < 0.001); one-way ANOVA with multiple comparisons in (A), (B); FDR calculated by rMATS program in (C), paired t-tests in (D)-(G), (I).

Given the strong correlation between HNRNPH1 and RBM3 expression, we speculated that HNRNPH1 expression is temperature dependent. To our surprise, HNRNPH1 protein levels or its interaction with spliceosomes, shown by core spliceosomal protein SmB pulldown, were not altered by cooling (Fig. S4E, S4F), suggesting that cooling-related regulation of RBM3 mRNA by HNRNPH1 is not driven by generic changes in HNRNPH1 spliceosomal association. We next explored whether the physical interaction between HNRNPH1 and RBM3 pre-mRNA is more important at 32°C by evaluating the efficiency of RBM3 PE removal when the HNRNPH1 binding site in the RBM3 intron is abolished. The intronic sequence between RBM3 Exon 3 and 4 contains multiple poly-G stretches (Fig. 4H), which are known as HNRNPH1 consensus binding sites (Caputi and Zahler, 2001; Uren et al., 2016). Indeed, analysis of published HNRNPH1 Individual-nucleotide resolution UV crosslinking and immunoprecipitation (iCLIP) dataset in HEK293T cells (Braun et al., 2018) revealed that HNRNPH1 strongly interacts with GGGG motifs at the 5’ end of PE (near 3’ splice site) and 30 nucleotides upstream of the 5’ splice site between RBM3 Exon 3 and 4 (Fig. 4H). Strikingly, the removal of this single GGGG motif was sufficient to dramatically increase PE inclusion and almost completely incapacitate its skipping upon cooling, but not at 37°C (Fig. 4I), recapitulating PE inclusion due to HNRNPH1 KD at 32°C (Fig. 4E). Taken together, the results show that HNRNPH1 interacts with the G-rich motif within the PE to repress RBM3 PE inclusion most efficiently upon cooling.

It is worth noting that G-rich sequences have a tendency to form RNA G-quadruplex (rG4) secondary structures (Kharel et al., 2020), which are highly enriched at weak splice sites and promote cassette exon inclusion (Georgakopoulos-Soares et al., 2022; Huang et al., 2017). Intriguingly, HNRNPH1 has been recently reported as an interactor and destabilizer of rG4s (Vo et al., 2022). Therefore, destabilising rG4s by HNRNPH1 is likely to induce exon skipping, as reported for HNRNPH1-mediated exon exclusion in RNA-binding protein EWS (EWSR1) transcripts (Vo et al., 2022). Consistent with this hypothesis, sequence-based rG4 prediction suggests the presence of rG4s in the region between RBM3 Exon 3 and 4 that overlaps with the primary HNRNPH1 binding site (Fig. 4H). Deletion of the GGGG motif in the mutant minigene is predicted to disrupt the rG4 (Fig. S4G), suggesting a potential connection between altered rG4 stability and temperature-sensitive alternative splicing of RBM3 mRNA.

## Discussion

Using a genome-wide CRISPR/Cas9 gene KO screen, we identified key trans-regulatory factors for neuronal RBM3 cold-induction. The strength of our screen lies in its closely recapitulating the physiological scenario, because we chose to a) GFP-tag endogenous RBM3 loci to account for any regulatory element beyond the coding sequence, b) use fully differentiated i-neurons with functional synapses, and c) pre-cool at 32°C to extend the dynamic range of GFP-RBM3 fluorescent readings affected by positive and negative regulator depletion. While splicing regulating proteins were identified among the strongest RBM3 regulators, our screen also indicated that RBM3 protein expression is controlled at multiple levels, from transcription, translation to protein degradation. It will be interesting to explore the overlap between cooling-dependent modulators of RBM3 identified in this study and regulators involved in changes of RBM3 levels by other known stimuli, such as BDNF/TrkB signalling cascade we previously reported (Peretti et al., 2021).

The appearance of HNRNPH1 as the top positive regulator of RBM3 together with other splicing factors prompted us to investigate cold-induced splicing changes in RBM3 mRNA. In line with this, we report cooling-dependent PE exclusion as a level of regulation governing RBM3 induction. Interestingly, this alternative splicing regulation is conserved in different cell types and between human and mouse (Preussner et al., 2022). Depletion of HNRNPH1 using different methods disrupted the alternative splicing control around RBM3 PE in endogenous RBM3 mRNA and in an externally introduced minigene. Moreover, removal of the poly-G stretches within the PE significantly repressed the removal of PE in RBM3 transcript, further conclusively establishing the role of HNRNPH1 binding.

It is interesting that depletion of HNRNPH1 leads to enhanced RBM3 PE retention at 32°C but exerted little impact at 37°C (Fig. 4D), supporting a cold-dependent efficacy of HNRNPH1 in RBM3 PE repression. The low efficiency of HNRNPH1 in RBM3 PE removal at 37°C is compensated by its overexpression (Fig. 4F and 4G), but this does not explain its cold-dependency, since HNRNPH1 expression is not changed upon cooling (Fig. S4E). Thus, the increased repressive capacity of HNRNPH1 may be explained by either or both of the following mechanisms. First, colder temperatures are known to enhance rG4s structural stability both directly (Moon et al., 2015; Yang et al., 2022) and indirectly due to modified enzymatic activities of RNA helicases and other RNA chaperones. Formation of rG4 could increase the capacity of HNRNPH1 to compete with other RBPs on the corresponding intronic region and thereby effectively repress the RBM3 PE. Alternatively, post-translational modification (PTM) and/or temperature-driven condensation of HNRNPH1 (Kim and Kwon, 2021) might modify its affinity for targeted RNA regions. Temperature variation can result in markedly different PTM patterns shaped by temperature-sensitive PTM enzymes (Cai et al., 2018), including kinases (Haltenhof et al., 2020) and arginine methyltransferases (Hong et al., 2010). In fact, in *vitro* and *in vivo* evidence supports temperature-dependent changes in RNA-RBP condensation (Iserman et al., 2020; Molliex et al., 2015; Pullara et al., 2022; Riback et al., 2017) as a result of differential intermolecular interactions (Tauber et al., 2020), possibly linked to distinct PTM of the embedded RBPs (Hofweber and Dormann, 2019; Sridharan et al., 2022).

Interestingly, HNRNPH1 missense mutations in the nuclear localisation sequence and nonsense mutations leading to reduced protein levels have been identified in individuals with neurodevelopmental disorders and intellectual impairment (Gillentine et al., 2021; Reichert et al., 2020). A possibility is that these mutations may act, in part, through impaired HNRNPH1 induction of RBM3 expression, affecting its roles in neurogenesis (Zhu et al., 2019) and/or synaptic maintenance (Peretti et al., 2015, 2021). In addition, HNRNPH1 sequestration in RNA foci has been implicated in neurodegenerative disease models of frontotemporal dementia, associated with impaired splicing functions (Bampton et al., 2020). It would be exciting to explore the effect of these HNRNPH1 mutations, and its sequestration in RNA foci, on RBM3 splicing and hence RBM3 protein levels.

The fate of inclusion or exclusion of alternative exons is a consequence of the interplay between the recruited splicing factors and the cis-acting elements with varying affinities, opening up opportunities for targeting different steps during mRNA splicing as therapeutic interventions (El Marabti and Abdel-Wahab, 2021). Although small molecules affecting global splicing activity has been applied to cancer treatments (Agrawal et al., 2018), modulating specific splicing events by targeting cis-acting elements using splice-switching antisense oligonucleotides (ASOs) has confirmed its therapeutic benefits in several genetic diseases, including spinal muscular atrophy and Duchenne muscular dystrophy (Havens and Hastings, 2016). Repressing PE inclusion in RBM3 using ASOs successfully upregulates RBM3 expression for neuroprotection in mice with prion neurodegeneration (Preussner et al., 2022). The past two decades have witnessed the development of splicing control using bifunctional oligonucleotides, consisting of an antisense domain complementary to the mRNA region close to the splice site, and a tail domain recruiting RBPs to promote exon inclusion or exclusion (Skordis et al., 2003; Zhou, 2022). Identifying HNRNPH1 as an RBM3 PE repressor brings the potential to apply this technology, providing novel targets to maintain RBM3 levels in both acute brain injury and in neurodegenerative disorder patients, with potential new therapeutic avenues for dementia treatments.

## Author Contributions

Conceptualization, J.Q.L., E.M., M.-D.R. and G.R.M.; Methodology, J.Q.L., D.K., and S.P.; Investigation: J.Q.L., D.K., S.P., S.M., S.F.F., J.M.Z. and J.L.F.; Formal analysis: S.M., N.P., F.C.Y.L. and J.K.; Supervision, J.U., E.M., M.-D.R. and G.R.M.; J.Q.L. and D.K. wrote the manuscript with inputs from other authors.

## Acknowledgements

The authors thank Sandeep Rajan and Stefanie Foskolou (University of Cambridge) for technical assistance, Oliver Muehlemann (University of Bern) for sharing the SMG1 inhibitor, Alessio Vagnoni (King’s College London) for HEK293T cells, Cambridge CRUK Core Genomics Facility and CIMR Flow Cytometry Core Facility for services. This work is supported by a Sir Henry Wellcome Postdoctoral Fellowship [215943/Z/19/Z] (to J.Q.L.), an Open Targets grant [OTAR2054] (to S.P., S.F.F., E.M., G.R.M.), the UK Dementia Research Institute (to D.K., S.M., N.P., J.U., E.M., M.-D.R., G.R.M.), which receives its funding from UK DRI Ltd, funded by the UK Medical Research Council (MRC), Alzheimer’s Society and Alzheimer’s Research UK. This research was funded in whole, or in part, by the Wellcome Trust and MRC. For the purpose of open access, the authors have applied a CC BY public copyright license to any Author Accepted Manuscript version arising from this submission.

## Declaration of interests

S.P. is now an AstraZeneca employee.

## Supplemental Information

**Figure S1.**
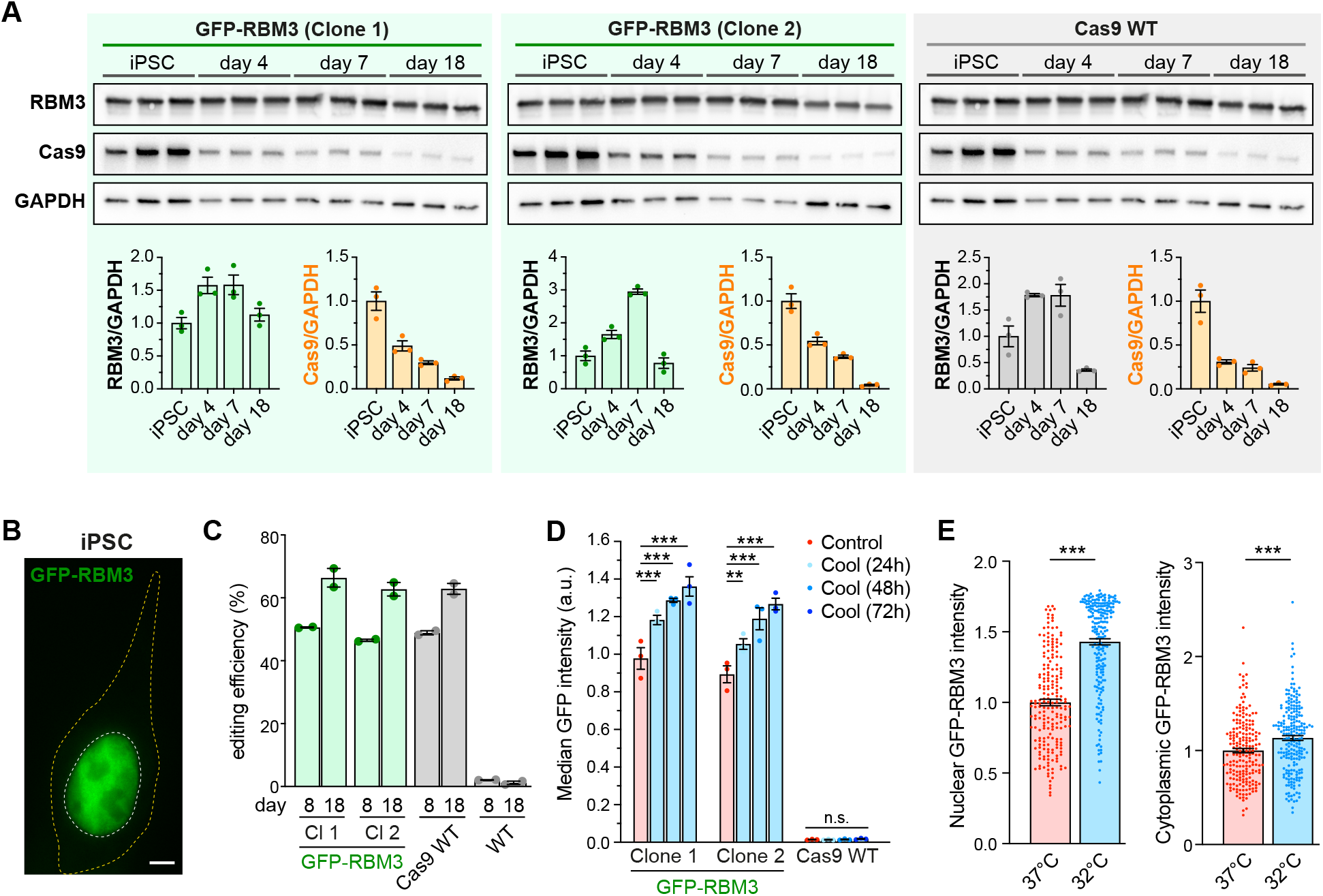
Characterisation of GFP-RBM3 human iPSC reporter line for CRISPR knockout screen. Related to Figure 1. (A) Western blots and quantification of RBM3, Cas9 and GAPDH in two GFP-RBM3 clones and Cas9 WT iPSCs and i-neurons. (B) Representative image of GFP-RBM3 iPSCs. The nucleus and soma are outlined by white and yellow dashed lines, respectively. (C) Editing efficiency of two GFP-RBM3 clones, Cas9 WT and WT i-neurons at day 8 and 18 post differentiation Median GFP intensity of two GFP-RBM3 clones and Cas9 WT i-neurons at 37°C or at 32°C for 24-72h, measured by flow cytometry. (D) Nuclear and cytoplasmic GFP intensity per unit area in GFP-RBM3 i-neurons at 37°C or 32°C (72h). N≥3 biological replicates. Mean ± SEM; **(p < 0.001); one-way ANOVA with multiple comparisons in (D), unpaired t-tests in (E). Scale bars: 5 μm.

**Figure S2.**
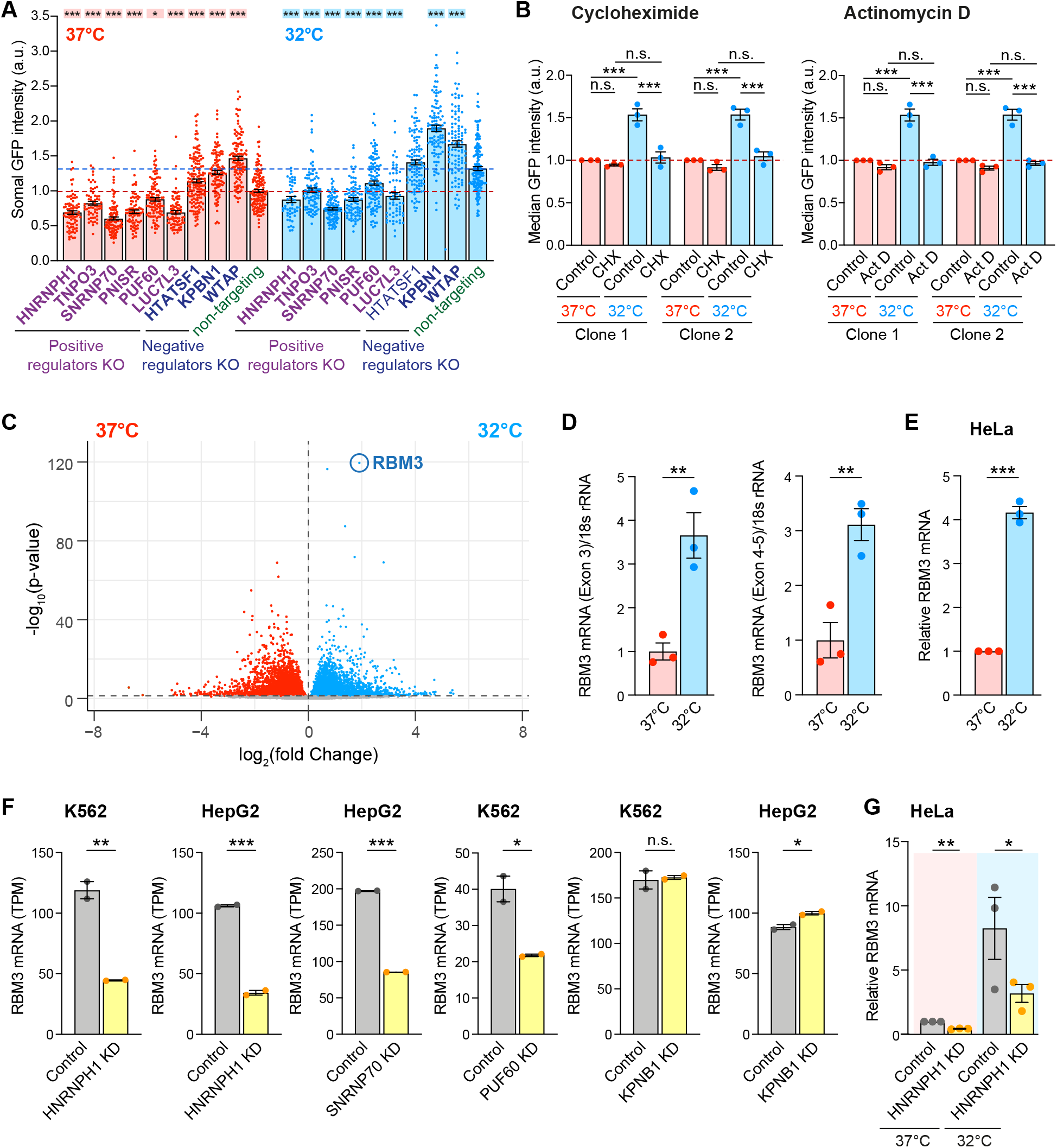
Cooling and regulators involved in mRNA splicing change RBM3 transcript levels. Related to Figure 2. (A) Somal GFP intensity per unit area in GFP-RBM3 i-neurons transduced with lentivirus containing specific or non-targeting sgRNA at 37°C or 32°C (72h) imaged by wide-field microscopy. Statistical analysis is performed between the specific and non-targeting sgRNA within the temperature groups. (B) Median GFP intensity per unit area of GFP-RBM3 i-neurons at 37°C or 32°C (72h) treated with cycloheximide (CHX) at 50 μM for 72h or actinomycin D (Act D) at 1 μM for 72h. (C) Volcano plot showing differential expression analysis of all transcripts identified in i-neurons at 37°C and 32°C (72h) from RNA-Seq data. (D) qRT-PCR of RBM3 Exon 3 or Exon 4-5 levels normalised to 18s rRNA in i-neurons at 37°C and 32°C (72h). (E) qRT-PCR analysis of RBM3 mRNA (Exon 2-3) normalised to GAPDH in HeLa cells at 37°C and 32°C (48h). (F) Normalised RBM3 mRNA abundance (TPM) of control and selective regulator candidates knocked-down K562 or HepG2 cells. Data are extracted from ENCODE project. 2 isogenic replicates are included in each condition. (G) qRT-PCR of RBM3 mRNA level normalised to GAPDH in control and HNRNPH1 KD HeLa cells at 37°C or 32°C (48h). N≥3 biological replicates. Mean ± SEM; n.s. (not significant), * (p<0.5), **(p<0.01), ***(p < 0.001); one-way ANOVA with multiple comparisons in (A), unpaired t-tests in (B), (D)-(F); paired t-test in (G).

**Figure S3.**
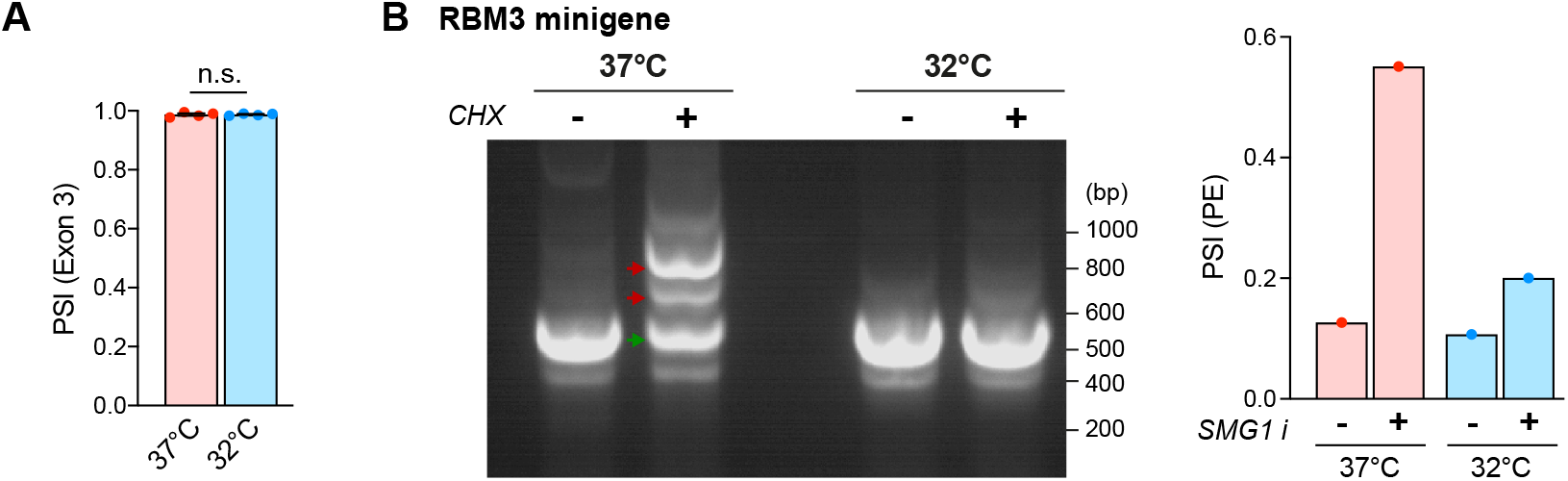
Cooling represses RBM3 poison exon inclusion. Related to Figure 3. (A) PSI values of RBM3 Exon 3 relative to Exon 2 and 4 in i-neurons at 37°C or 32°C (72h). N=4. (B) RT-PCR of RBM3 minigene expressed in HeLa cells at 37°C and 32°C (48h) in the presence or absence of cycloheximide (CHX)(200ug/ml). PSI values of RBM3 PE are shown in the graph. N=1. Mean ± SEM; * (p<0.5), **(p < 0.01), ***(p < 0.001); FDR calculated by rMATS program in (A).

**Figure S4.**
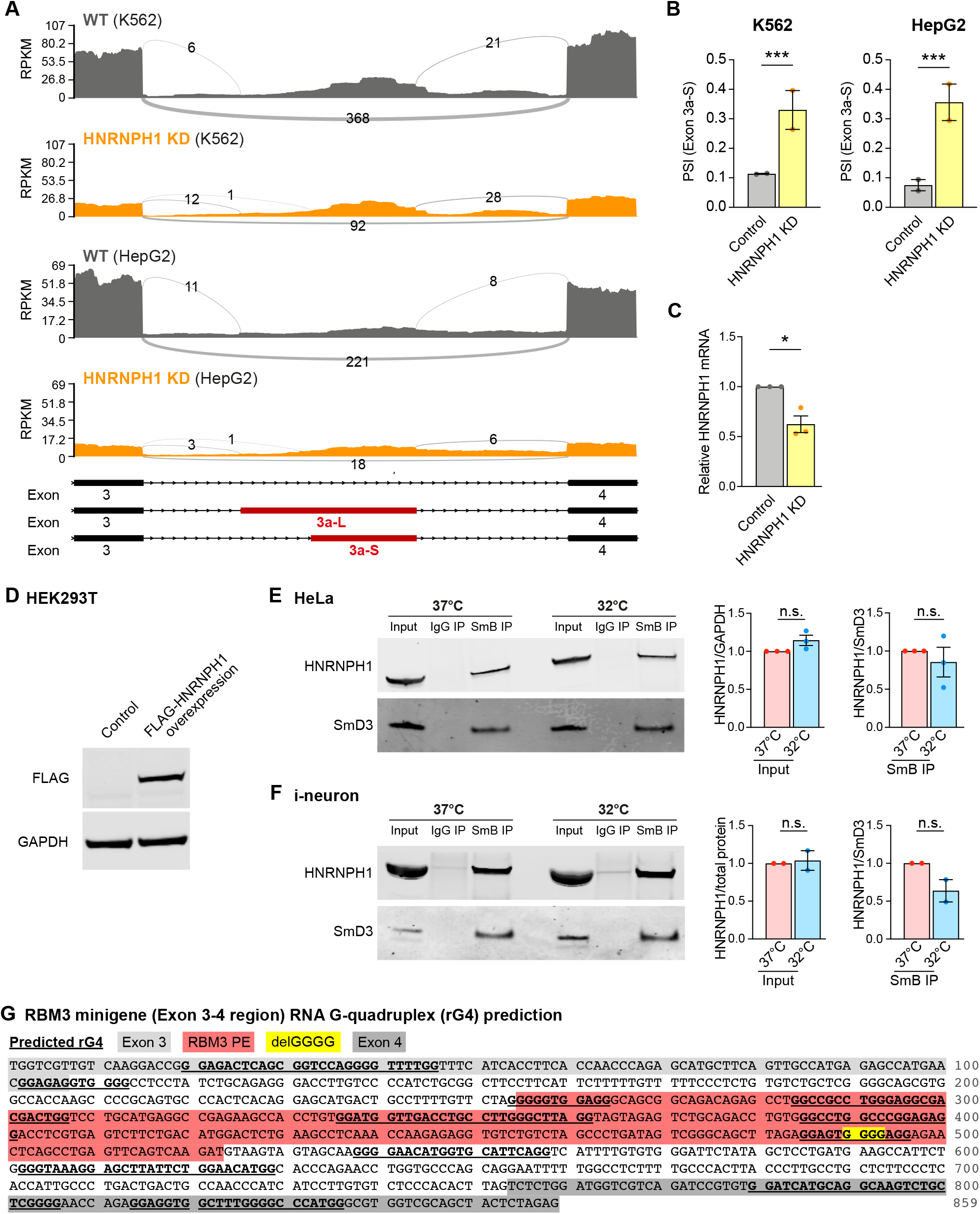
HNRNPH1 expression enhances RBM3 mRNA poison exon skipping on cooling. Related to Figure 4. (A) Sashimi plots of the region between Exon 3 and 4 of RBM3 transcripts in control and HNRNPH1 knocked-down K562 and HepG2 cells, showing major alternatively spliced isoforms. Data are from ENCODE Project. 2 isogenic replicates are included. (B) PSI values of RBM3 Exon 3a-S in control and HNRNPH1-knocked down K562 and HepG2 cells. RNA-Seq data from ENCODE Project, 2 isogenic replicates are included. (C) qRT-PCR of HNRNPH1 mRNA normalised to GAPDH upon HNRNPH1 KD in HeLa cells. (D) Western blots of FLAG-HNRNPH1 and GAPDH (loading control) in control and FLAG-HNRNPH1-overexpressed HEK293T cells at 37°C. (E) Western blot and quantification of HNRNPH1 total protein levels (input HNRNPH1 normalised to GAPDH) and its abundance in spliceosomal protein SmB pulldown (normalised to SmD3) in HeLa cells at 37°C and 32°C (48h). (F) Western blot and quantification of HNRNPH1 total protein levels (input HNRNPH1 normalised to ponceau measured total protein abundance) and its abundance in spliceosomal protein SmB pulldown (normalised to SmD3) in i-neurons at 37°C and 32°C (72h). (G) RNA G quadruplexes (rG4) within the RBM3 Exon 3-4 region are predicted using QGRS mapper. Deletion of the GGGG motif in the mutant RBM3 minigene is predicted to disrupt the rG4 structure overlapping this region. N≥3. Mean ± SEM; n.s. (not significant); FDR calculated by rMATS program in (B), paired t-tests in (C)-(F).

## Key resources table

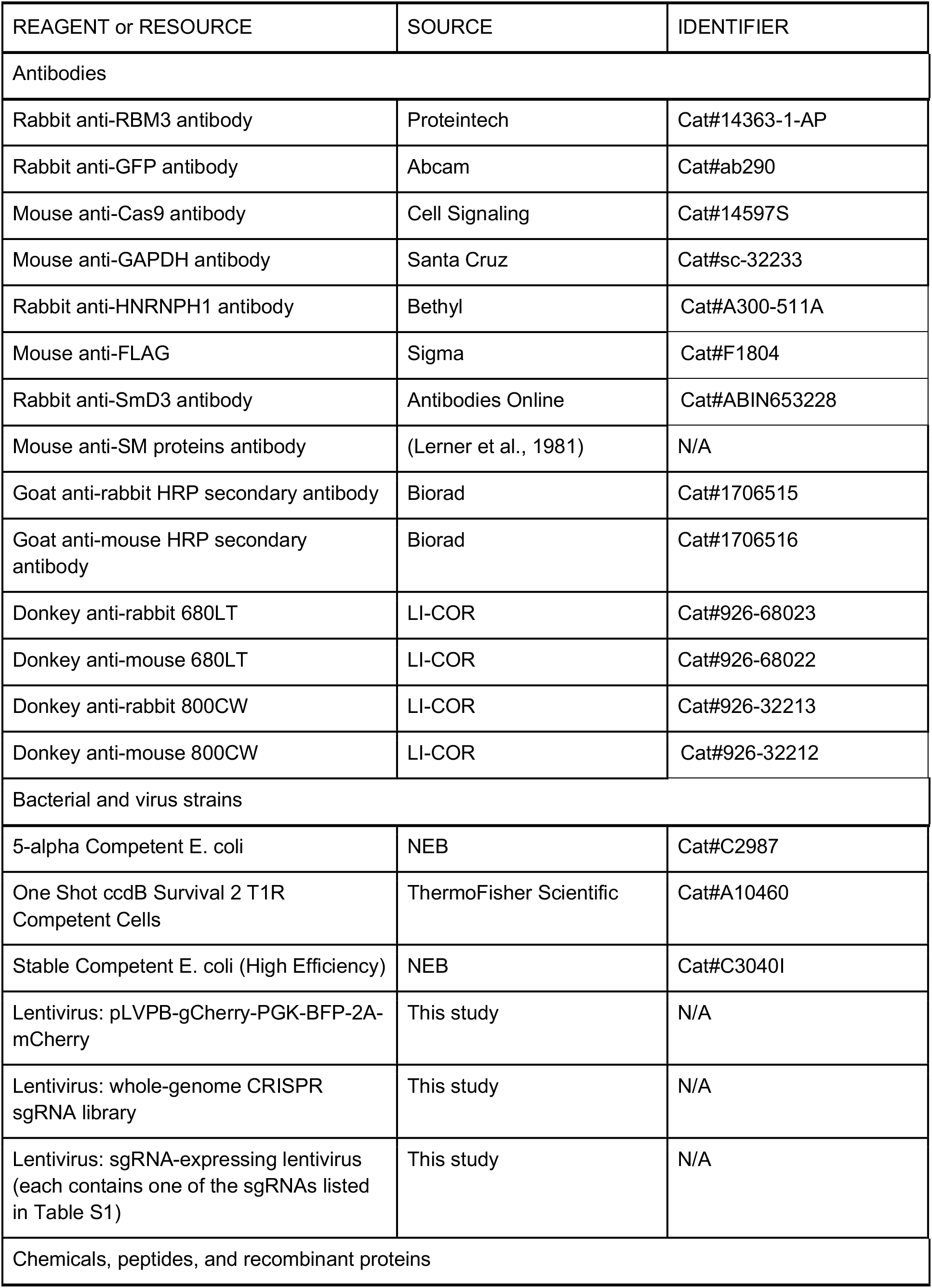
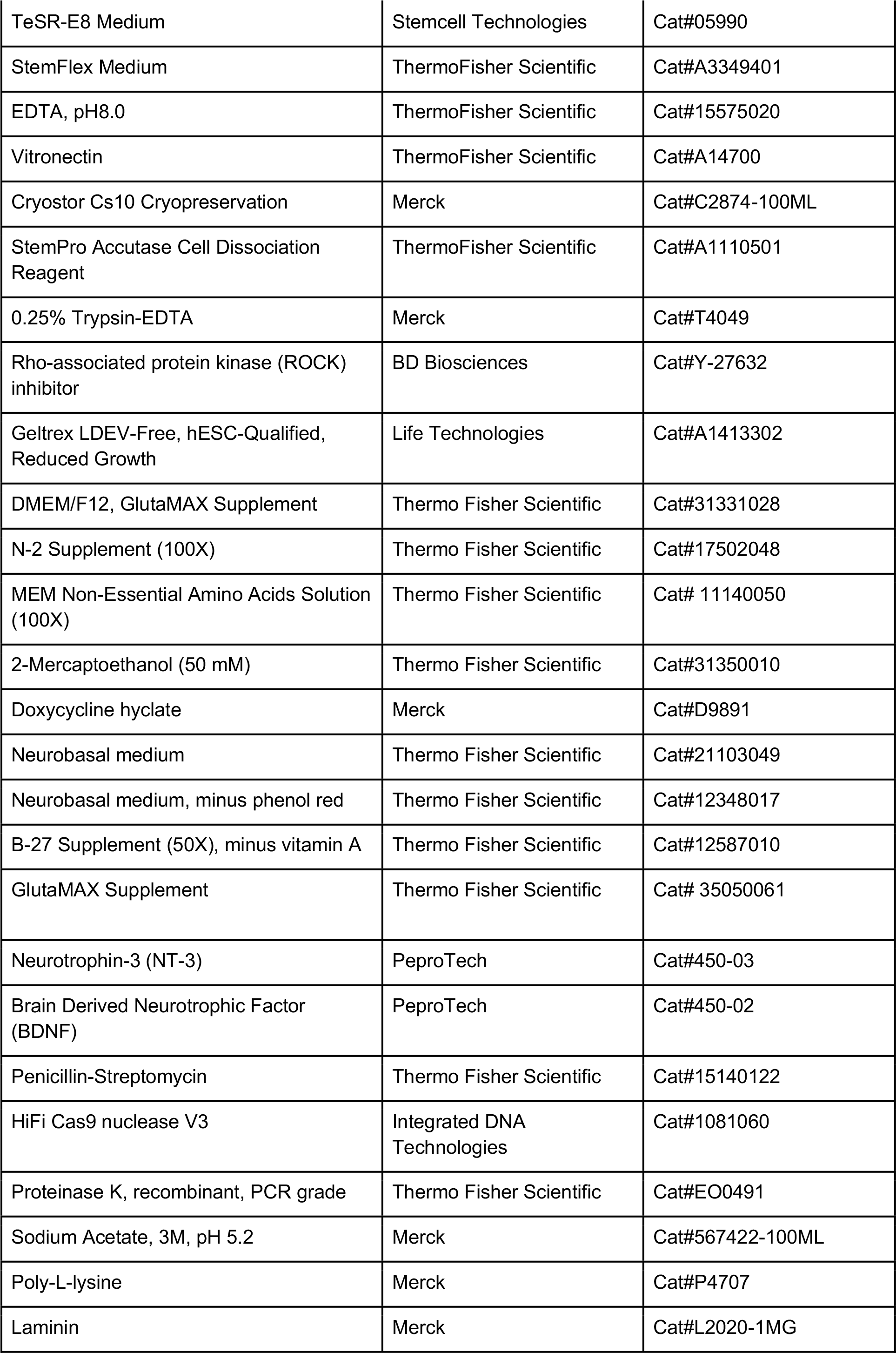
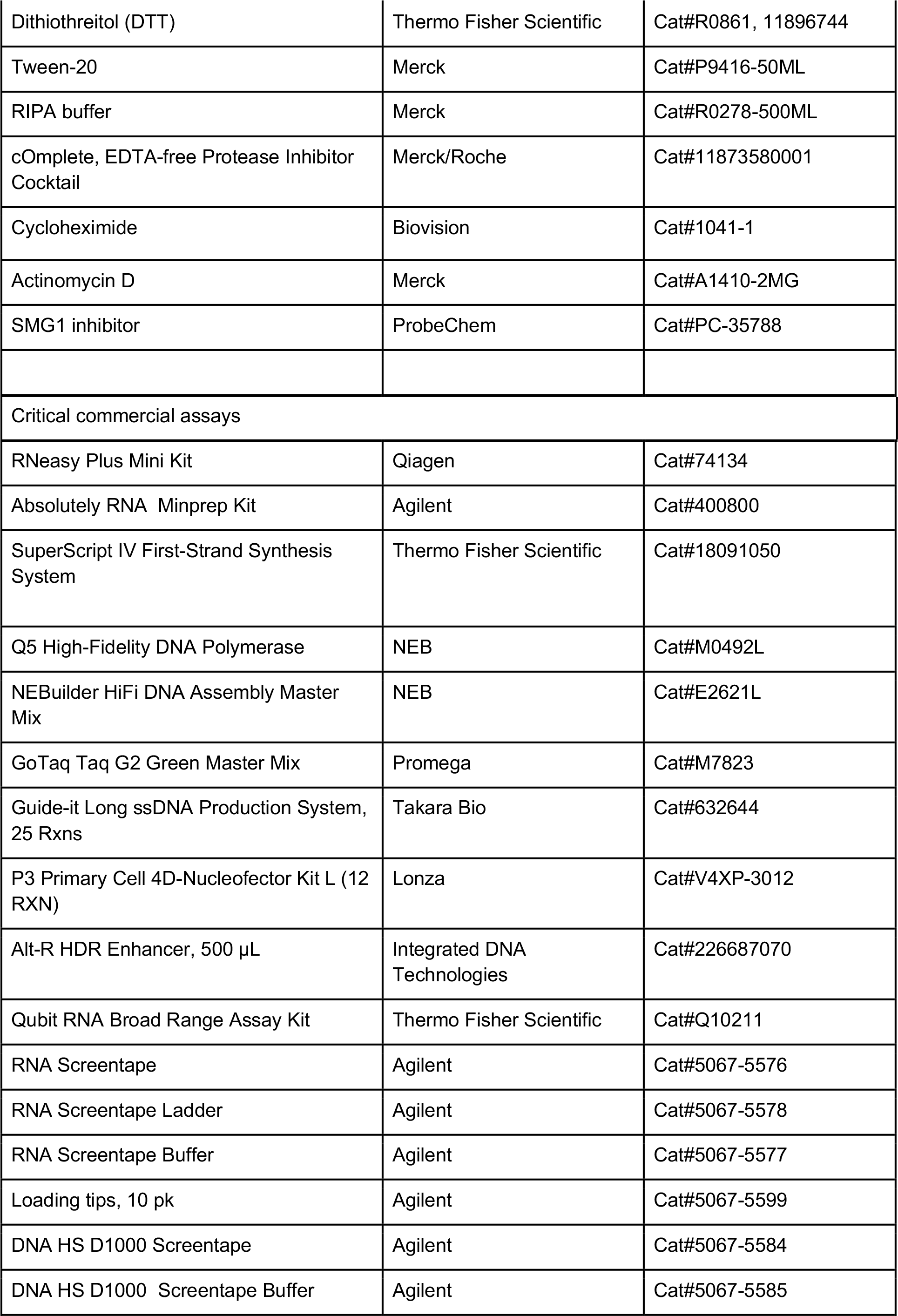
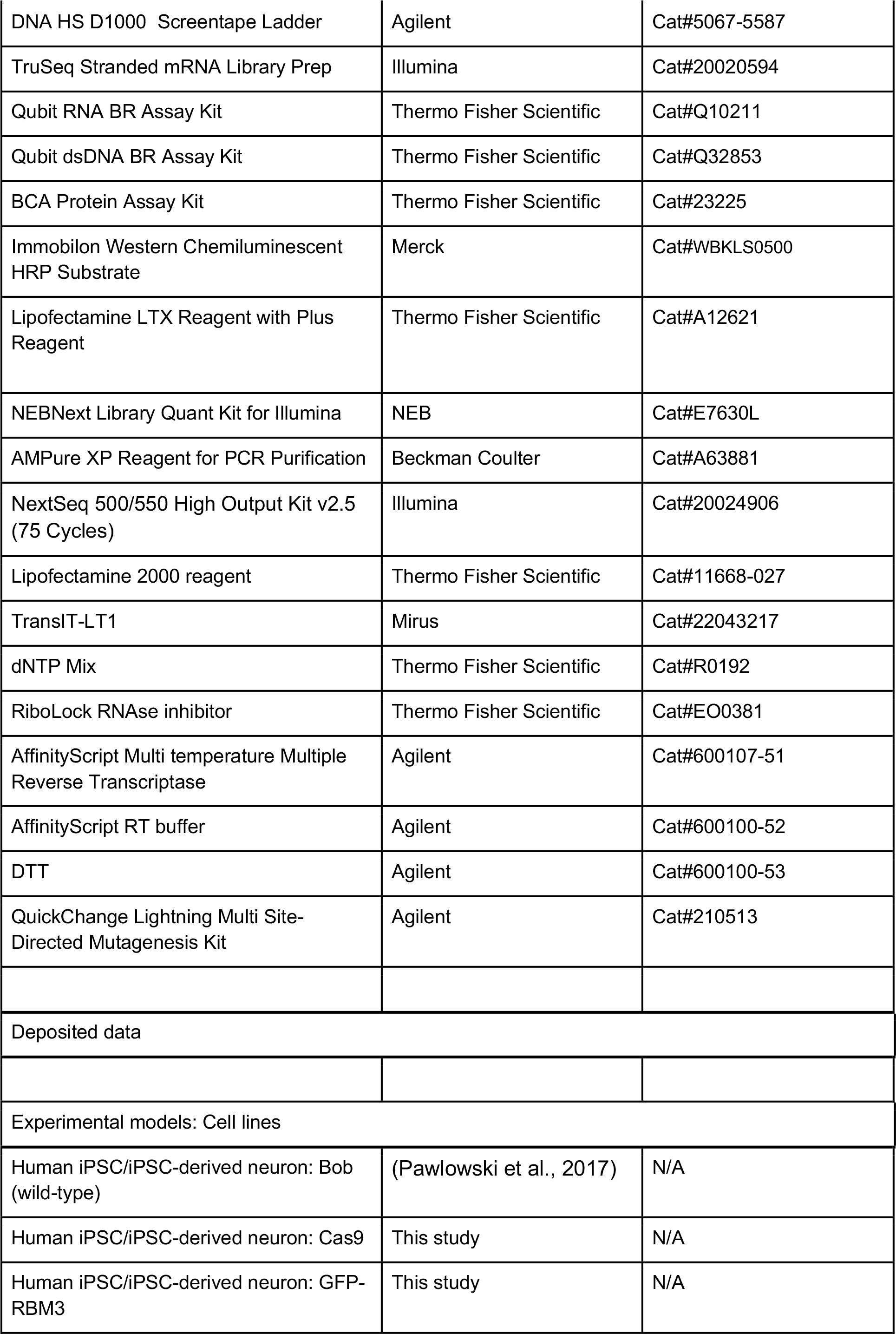
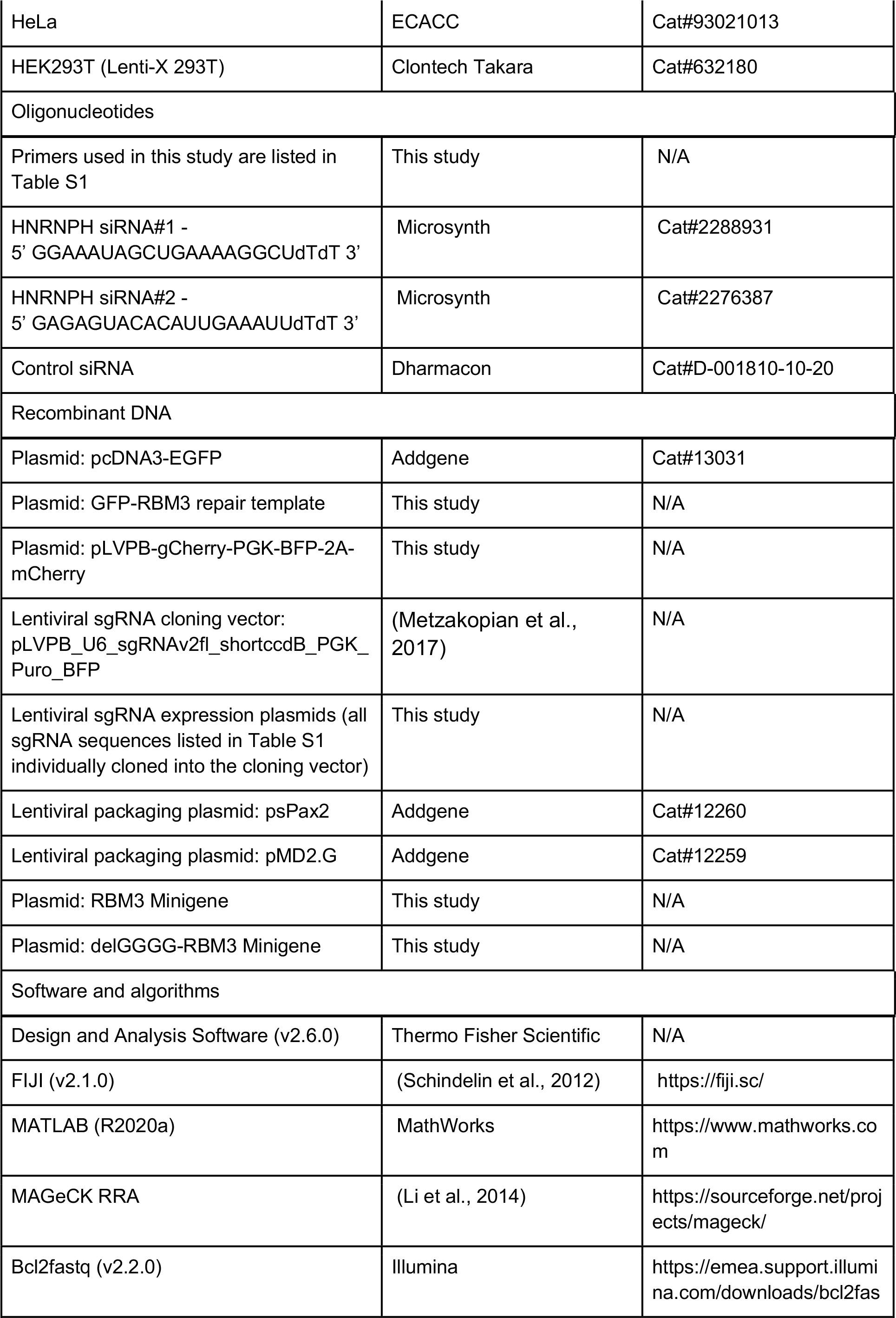
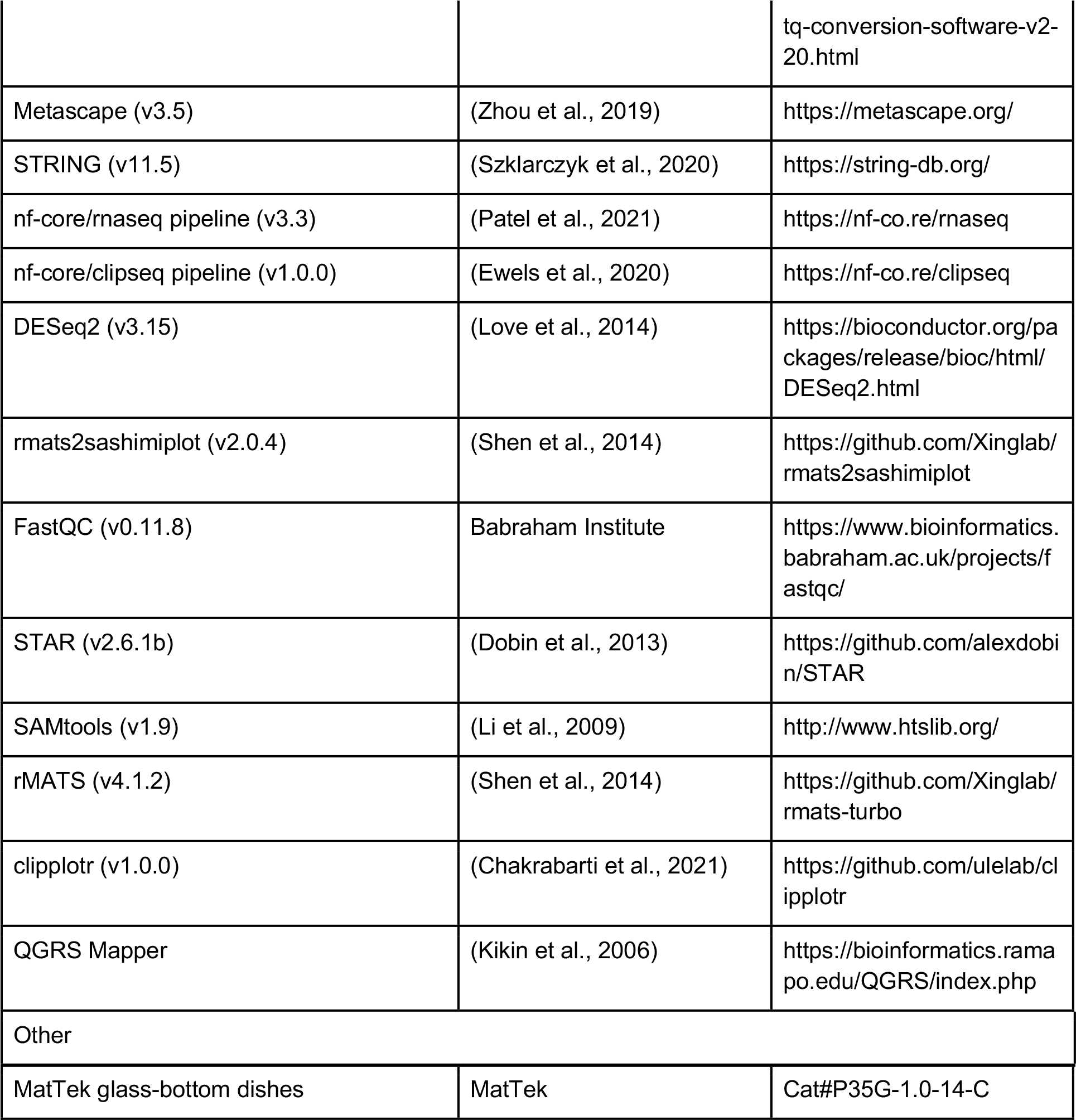

## Human iPSC culture

Human induced pluripotent stem cells (iPSCs) with Neurogenin-2 (NGN2) transgene stably integrated into a ‘safe-harbour’ locus under doxycycline (Dox)-inducible promoter (Pawlowski et al., 2017) were maintained under feeder-free conditions in a 37°C, 5% CO_2_ tissue culture incubator. They were cultured on vitronectin (3.3 μg/mL)-coated culture plates or glass-bottom dishes and fed every day with TeSR-E8 medium or every 2 days with StemFlex Medium. 0.5mM EDTA was used for routine dissociation to maintain colony growth. iPSCs were frozen in Cryostor Cs10 Cryopreservation.

### iPSCs differentiation into iPSC-derived neurons (i-neurons)

iPSCs were enzymatically detached and dissociated into single cells using Accutase and plated into GelTrex (1:100 dilution)-coated culture plates in TeSR-E8 medium supplemented with 10 μM Rho-associated protein kinase (ROCK) inhibitor. After 24 hours (day 1), TeSR-E8 medium was changed to DMEM/F12 medium with GlutaMAX, supplemented with 1x N-2 supplement, 1x Non-Essential Amino Acids, 50 nM 2-Mercaptoethanol, 100 U/mL Penicillin-Streptomycin, and 1 μg/mL Doxycycline Hyclate (Dox) (iN-1 medium). After 24 hours (day 2), the medium was replaced with the same medium as the previous day. From day 3 to day 6, the culture was fed daily with Neurobasal medium supplemented with 1x B-27 supplement (minus vitamin A), 1x GlutaMAX, 50 nM 2-Mercaptoethanol, 100 U/mL Penicillin-Streptomycin, and 1 μg/mL Dox, 10 ng/mL Neurotrophin-3 (NT-3), and 10 ng/mL Brain-derived neurotrophic factor (BDNF) (iN-2 medium). After day 6, the medium was changed every other day.

Cultures used for flow cytometry were prepared by dissociating day 4 iPSCs with Accutase and plating 80,000 cells per well in a 96-well culture plate pre-coated with GelTrex (1:100 dilution). The same feeding schedule as previously described was followed from day 5.

To prepare cultures for live fluorescent imaging, day 4 iPSCs were dissociated with Accutase and plated 10,000-20,000 cells/dish on 35 mm MatTek glass-bottom dishes pre-coated with 0.1 mg/mL poly-L-lysine and 10 μg/mL laminin in iN-2 medium with ROCK inhibitor. The same feeding schedule as previously described was followed from day 5.

## HeLa and HEK293T cell cultures and transfection

HeLa and HEK 293T cells were grown in Dulbecco’s modified Eagle’s medium (DMEM)+ Ham’s F12 (1:1) supplemented with 10% fetal calf serum (FCS), penicillin (100 IU/mL), and streptomycin (100 μg/mL) and grown at 37°C and 5% CO2. Plasmid DNA transfections were performed with Lipofectamine 2000 (Thermo Fisher Scientific) and TransIT-LT1 (Mirus) according to the manufacturer’s instructions. For HNRNPH1 siRNA transfections, 250,000 HeLa cells were seeded in a 6-well plate and the next day cells were transfected with two siRNAs 10nM siRNA each (siRNA #1 + siRNA#2) using Lipofectamine 2000. After 24h, a second round of siRNA transfection is done in fresh medium with or without RBM3 minigene co-transfection. One set of the cells is moved to 32°C while the other set is kept at 37°C. 24h later, Medium is changed again with fresh medium with or without SMG1 inhibitor (1uM). After 24h, cells are harvested. SMG1 inhibitor treatment (1uM) and Cycloheximide (CHX) treatment (200ug/ml) was done for 24h before harvesting the cells.

## Plasmids, oligonucleotides, and guide RNAs

Plasmids: RBM3 minigene was cloned by amplifying genomic regions from exon 1-4 using forward primer - AAGAATTCATGTCCTCTGAAGAAGGAAAGC and reverse primer – TTTGCGGCCGCCTCTAGAGTAGCTGCGACCACGCC and then inserting in EcoRI-NotI sites in pcDNA3.1(+) vector backbone. Minigene Del GGGG mutant was generated using site directed mutagenesis (QuickChange Lightning Multi Site-Directed Mutagenesis Kit) using the primer-GTCGGGCAGCTTAGAGGAGTAGGAGAACTCAG. FLAG HNRNPH1 WT expression construct was generated by inserting HNRNPH1 coding sequence, obtained as a string synthesised (GeneArt, Thermo Fisher Scientific), in XbaI-NotI sites in pcDNA6F vector backbone. All primers, siRNAs and guide RNAs (sgRNAs) used in this study are listed in Table S1.

To prepare iPSC cDNA, RNA was purified from iPSC pellets using the RNeasy Mini kit and reverse transcribed into cDNA using the oligo(dT) primer and the SuperScript IV First-Strand Synthesis System. The following fragments were amplified by PCR using Q5 DNA Polymerase and the indicated templates and primers (see Table S1): a) The 5’ homology arm (1 kb sequence immediately upstream of RBM3 start codon) was amplified from iPSC cDNA using Primers Pr1 and Pr2; b)The GFP fragment was amplified from pcDNA3-EGFP using Primers Pr3 and Pr4; c) The 3’ homology arm (start codon and 1 kb sequence at it immediate downstream) was amplified from iPSC cDNA using Primers Pr5 and Pr6; d) the origin of replication and ampicillin-resistant gene were amplified using Primers Pr7 and Pr8.

The repair template plasmid for CRISPR knocked in of GFP to the N-terminal of the RBM3 coding region (GFP-RBM3 repair template) was generated by assembling Fragments a-b-c-d in the indicated order using NEBuilder. Plasmids were validated by Sanger sequencing. Single-stranded DNA (ssDNA) were subsequently synthesized from these repair template plasmids to improve CRISPR knock-in efficiency using Guide-it Long ssDNA Production System with Primers Pr9 and Pr10 according to manufacturer instructions.

The 19-nucleotide (nt) guide RNA (gRNA) RBM3-N gRNA#1 (5’-CUGCCAUGUCCUCUGAAGA-3’) and #2 (5’-UUUCCUUCUUCAGAGGACA-3’) followed by the protospacer adjacent motif (PAM) targeting the region adjacent to the RBM3 start codon were resuspended in water to the concentration of 4 μg/μL.

### Generation of GFP-RBM3 iPSCs by CRISPR

Half a million wild-type iPSCs were electroporated with Cas9 protein, RBM3 gRNAs and GFP-RBM3 repair template using Lonza Nucleofector Technology according to manufacturer instructions. Briefly, Cas9-RNP complex mixture containing 0.2 μL 3 M NaCl, 1 μL RBM3-N gRNA#1, 1 μL RBM3-N gRNA#2, 1 μL Cas9 protein (HiFi Cas9 nuclease V3, 10 μg/μL) were assembled and incubated at room temperature for 45 min. 0.5 million iPSCs were resuspended with the nucleofector solution (90 μL P3 solution and 20 μL supplement). 3 μL of the repair template ssDNA (4 μg/μL) were added to the pre-assembled Cas9-RNP, mixed with the iPSC suspension, and transferred to the Nucleocuvette Vessels. Placed the vessel into the nucleofector unit and started the program to complete electroporation. Immediately after, the electroporated cells were gently transferred to the 4 mL pre-warmed StemFlex medium supplemented with 10 μM ROCK inhibitor and 40 μL homology-directed repair (HDR) enhancer and plated into 2 vitronectin-coated wells in a 6-well plate. The cells were then incubated at 32 °C for 48 h. The following day after electroporation, the medium was replaced with StemFlex medium and replaced every other day. When reaching 70-80% confluency, cells in one well were frozen in Cryostor Cs10 Cryopreservation at −80 °C and those in the other were detached and dissociated with Accutase. Isolated iPSCs were plated in a vitronectin-coated 96-well plate in StemFlex medium supplemented with 10 μM ROCK inhibitor, with 30-50 cells per well. From the next day, the iPSCs were fed with StemFlex medium every other day until confluent. Confluent wells were dissociated and half of the cells were subject to flow cytometry measurement (see Flow cytometry) to determine the GFP intensity of iPSCs in each well. 500-750 cells from each of the wells with GFP-positive cells were plated in a vitronectin-coated 10 cm petri dish in StemFlex medium supplemented with 10 μM ROCK inhibitor. From the next day, the hiPSCs were fed with StemFlex medium every other day until colonies were formed (1-2 weeks). Imaged with a wild-field fluorescent microscope, GFP-positive colonies were picked by gentle aspirating with P1000 pipette tips and transferred to a well in round-bottom 96-well plates with 200 μL StemFlex medium with 10 μM ROCK inhibitor and 100 U/mL Penicillin-Streptomycin per well. Once finished colony picking, each well was split into 2 vitronectin-coated flat-bottom 96-well plates by gentle pipetting to break colonies into small clusters and transferring 100 μL cell suspension in each well into a well in new plates. Each well was fed StemFlex medium every other day until the majority of the wells reached 50-60% confluency. Genomic DNA of individual colonies in one of the duplicated plates was extracted (see Genomic DNA extraction) and PCR-genotyped using GoTaq Taq G2 Green Master Mix and primer pairs Pr11/12, Pr11/14, and Pr12/13. The correctly genotyped clones in the corresponding second plates were expanded in 6-well plates to be further validated by Sanger sequencing. Sequence-verified clones were aliquoted and frozen at −80 °C.

## Genomic DNA extraction

iPSCs were detached using Accutase and pelleted by centrifugation at 250 g for 5 min. After removing the supernatant, 50 μL (each well of a 96-well plate) or 5-10 μL (every 1000 iPSCs) lysis buffer (and 1:40 freshly added Proteinase K) was added to the pellet and lysed at 55 °C overnight in a shaker. The next day, 10% volume of 3M sodium acetate (pH 5.2) and an equal volume of isopropanol were added to the lysate and mixed by brief vortexing. Genomic DNA was pelleted by 15 min centrifugation at maximum speed, followed by two washes with 200-1000 μL of 80% ethanol. After the last centrifugation to remove the residual ethanol, pellets were left at room temperature to air dry, and resuspended in 50-500 μL TE buffer (10 mM Tris-HCl, pH 8.0 and 0.1 mM EDTA).

## Western blotting

Protein concentrations of lysates were determined by BCA assay following the manufacturer’s instruction. Samples were diluted with Laemmli protein sample buffer with 100 mM DTT. 10-15 μg protein was loaded into each well in pre-cast 12% gels or 4-15% gradient gels and ran at 125 V. Proteins were transferred to a 0.2 μm nitrocellulose membrane at 70 V for 2 h in a wet blot system. Membranes were blocked with 5% BSA or 5% non-fat milk in 1x TBS-T for 1 h rotating at room temperature. The primary antibody solution was incubated overnight at 4°C while rotating. The next day, membranes were washed three times with 1x TBS-T, then incubated for 1 h in secondary antibody solution (1:10,000 in 5% BSA or 5% non-fat milk in 1x TBS-T) and washed three times with 1x TBS-T before adding HRP chemiluminescent substrate to develop on a ChemiDox MP Imaging System or directly on LI-COR Odyssey CLx Primary and secondary antibodies were used in the following concentrations: rabbit anti-RBM3 (1:1000), mouse anti-Cas9 (1:1000), mouse anti-GAPDH (1:2000), mouse anti-Cas9 (1:2000), rabbit anti-HNRNPH1 (1:1000), mouse anti-FLAG (1:2000), rabbit anti-SmD3 (1:1000), donkey anti-rabbit 680LT (1:10000), donkey anti-mouse 680LT (1:10000), donkey anti-rabbit 800CW (1:10000), donkey anti-mouse 800CW (1:10000), goat anti-rabbit and goat anti-mouse HRP conjugated secondary antibodies (1:10000, Biorad).

## Lentiviral production

96-well plates or T75 flasks pre-coated with 25 μg/ml PLL at 37°C overnight, followed by 3 times washes with distilled water and left to dry in tissue culture hoods. HEK293FT were plated at 10^6^ cells/cm^2^ density and incubated at 37°C overnight. Lentiviral expression and packaging plasmids were transfected using Lipofectamine LTX. For each well of a 96-well plate (upscale proportionally for T75 flask transfection), Mix A: 20 μL Opti-MEM, 19.12 ng psPax2, 12.5 ng pMD2.G, 25 ng expression plasmid, 0.1 μL Plus Reagent; Mix B: 5 μL Opti-MEM, 0.3 μL Lipofectamine LTX were assembled and incubated at room temperature for 20 min before adding to the HEK293FT cultures. After 24h, replace with fresh media and check for BFP expression. Viruses were harvested 72h post-transfection by centrifuging at 6000g at 4°C overnight. Resuspend pellets with PBS and freeze at - 80°C for storage.

## Flow cytometry

Dissociated cells were resuspended in culture media in 96-well plates and fluorescent intensities of GFP, mCherry and/or BFP were measured at room temperature with a CytoFLEX S System (Beckman Coulter) and CytExpert (v2.4) Program. Acquired data were analysed with FlowJo (v10.7.2).

## Imaging of i-neurons

I-neurons cultured at low density on 35mm glass bottom dishes were imaged in phenol red-free iN-2 media (minimise autofluorescence) in OkoLab temperature-controlled chamber. Cells were imaged at two temperatures: 37°C (control and rewarmed) and 32°C (cooled) with 5% CO_2_. Cells were imaged with a custom-built wide-field microscope in Epi fluorescence mode with a 100x oil TIRF objective (Olympus). BFP was excited with a 504 nm laser at 10 mW, and GFP with a 488 nm laser at 5 mW. Both channels were imaged simultaneously and separated with OptoSplit III System (Cairn Research) and a Prime BSI camera (Photometrics).

## RBM3 CRISPR knockout screen

Both clones of GFP-RBM3 iPSCs were differentiated in GelTrex-coated T75 flasks. The total number of i-neurons used for each clone in each replicate was approximately 60,000,000 to ensure sufficient copies of individual sgRNAs could be obtained from the pool. 4 days post differentiation, the cultures were transduced with a whole-genome sgRNA lentiviral library (399 non-targeting sgRNAs and 91138 sgRNAs targeting 18466 genes across the human genome), co-expressing a BFP fluorescent reporter to indicate successfully transduced cells. The viral titer was determined in pilot experiments, resulting in 20% of the i-neurons becoming BFP-positive on day 18 to minimise multiple sgRNA entries into a single cell. I-neurons at 14, 15 or 16 days post differentiation (or 11, 12, 13 days post-transduction) were cooled at 32°C for 72h before dissociation with 20 min Accutase and 10 min Trypsin-EDTA incubation, followed by fluorescence-activated cell sorting (FACS). GFP intensity of all BFP-positive cells was manually separated into 4 quartiles, and only cells with the GFP intensity falling in the top or the bottom quartile were collected in separate tubes and their genomic DNA was extracted (see Genomic DNA extraction) immediately after. Around 3,000,000 i-neurons were collected in the low GFP or high GFP tube for each clone in each replicate.

Purified DNA was sequenced to identify sgRNAs enriched in high or low GFP populations. The sequencing library was created in a two-stage PCR reaction. The first stage amplifies the enriched gRNA cassettes: 2 μg DNA, 1.5 μL Forward Primer (10 μM), 1.5 μL Reverse Primer (10 μM), 25 μL NEB Q5 High-Fidelity 2x Master Mix in a 50 μL reaction. The PCR condition was: 98C for 30 sec, 25 cycles of (98C for 10 sec, 62C for 30 sec, 72C for 15 sec), and 72C for 2 min. PCR product was purified using Ampure XP Beckman magnetic beads and diluted to 200 pg/μL. The second PCR attached the Illumina adaptors and barcodes: 1 μL PCR product from the first reaction, 0.75 μL Forwards Primer (20 uM), 0.75 μL Reverse Primer (20 uM), 10 μL Roche 2X KAPA HiFi ReadyMix in a 20 μL reaction. The PCR condition was: 95C for 3 min, 9 cycles of (98C for 20 sec, 66C for 15 sec, 72C for 20 sec), and 72C for 1 min. PCR products were purified using Ampure XP Beckman magnetic beads and eluted in 35 μL TE buffer. PCR products were quantified with NEBNext Library Quant Kit for Illumina. The library was run on an Illumina NextSeq 550, using an Illumina 75 cycle, high output kit at 1.4 pM.

## Arrayed CRISPR target validation

Inserts containing the candidate gene targeting sgRNA (1-3 sgRNAs per gene) sequences and non-targeting sgRNA sequences (see Table S1) were cloned into lentiviral expression vector pLVPB-U6-sgRNAv2fl_shortccdB_PGK_Puro_BFP linearised with BbsI, which removed the suicide ccdB cassette. The sgRNA expression plasmids were verified by Sanger sequencing and used to generate lentiviral particles (see Lentiviral production) in a 96-well array format. GFP-RBM3 Clone 1 iPSCs at 4 days after differentiation in 96-well plates were transduced with the arrayed lentiviral library at a viral concentration pre-determined in pilot experiments to obtain maximal transduction efficiency. 15 days post differentiation, GFP-RBM3 i-neurons were either cooled at 32°C for 72h or continue to grow at 37°C. On day 18, BFP and GFP intensity of the transduced cultures were measured using flow cytometry (see Flow cytometry).

## RNA-Seq

Four biological replicates of WT i-neurons in control (37 °C) and cooled (72 h at 32 °C, day 15-18) were included in this study. Total RNA was extracted with RNeasy Plus Mini Kit. RNA concentrations of individual samples were measured using Qubit RNA Broad Range Assay Kit and their integrity was determined using Agilent TapeStation System. The library was prepared using Illumina TruSeq Stranded mRNA Library Prep following the manufacturer’s instructions and sequenced paired-end 150 bp on Novaseq.

## RT-PCR and qRT-PCR

RNA of i-neurons, HeLa and HEK293T cells in 3 or more biological replicates was extracted using the RNeasy Plus Mini kit or Absolute RNA miniprep kit and reverse transcribed into cDNA using random hexamers and the SuperScript IV First-Strand Synthesis System or AffinityScript Multiple Temperature Reverse Transcriptase. Briefly, to reverse transcribe RNA into cDNA, 1-2 μg of isolated RNA were incubated with random hexamers (900ng) in a total volume of 67 μl in DEPC water at 65 °C for 5 minutes and then for 10 minutes at room temperature. Then, the mixtures were separated into 2 parts – RT and no-RT control samples (33.5 μl each). To this 13.5 μl master mix were added to supplement 1x reverse transcriptase buffer (Agilent), 10 mM DTT (Agilent), 400 μM dNTPs (Thermo Scientific), 40 units RiboLock RNase inhibitor (Thermo Scientific) and 1 reaction equivalent AffinityScript Multiple Temperature Reverse Transcriptase (Agilent). The RT enzyme was absent in the no-RT condition. The samples were then incubated at 65 °C for 1 hour followed by heat-inactivation at 75 °C for 20 minutes. For RT-PCR, 20-40ng cDNA was amplified using GoTaq Polymerase mix using either RBM3 Exon 2-4 primers or RBM3 minigene specific primers (See also Table S1). Quantification of the bands on agarose gels was done using FIJI software. Percent spliced in (PSI) indexes of RBM3 poison exon (PE) are calculated based on the intensity of PE-included (red arrows) and PE-skipped (green arrows) isoforms visualised in agarose gels:

a. One PE-included isoform is detected: (I_Inc_/L_Inc_) / (I_Inc_/L_Inc_ + I_skip_/L_skip_), or
b. Two PE-included isoforms are detected: (I_Inc1_/L_Inc1_ + I_Inc2_/L_Inc2_) / (I_Inc1_/L_Inc1_ + I_Inc2_/L_Inc2_ + I_skip_/L_skip_)

Where I_Inc_ = Intensity of PE-included isoform, L_Inc_ = length of PE-included isoform in base pairs, I_Inc_ = Intensity of PE-skipped isoform, L_skip_ = length of PE-skipped isoform in base pairs.

For qRT-PCR, cDNA of each sample (diluted 10-1000 times) was mixed with SYBR Green PCR Master Mix and PCR primers in 4 technical replicates in 384-well plates. Reactions were performed in a QuantStudio Real-Time PCR system (ThermoFisher Scientific). Analysis was performed with Design and Analysis Software (v2.6.0, ThermoFisher Scientific). PSI indexes of RBM3 PE are ratios of the normalised expression levels between RBM3 Exon 3a amplicon and RBM3 Exon 3 (or Exon 4-5) amplicon.

## Data analysis

### Image quantification of GFP-RBM3 iPSCs and i-neurons

Regions with single cells were cropped from the full-size images with a custom-made program. Then, each single cell image was analysed with a custom-made MatLab pipeline. Briefly, two channels were used for analysis: BFP and GFP. BFP is expressed by the lentivirus as a reporter to indicate transduced cells and it is used to define the cell boundary. Nuclei were defined as areas where GFP is uniformly expressed at higher intensities, as RBM3 is predominantly nuclear. This was achieved by applying automatic global Otsu’s threshold to the GFP channel. Average GFP channel intensities (total intensities divided by corresponding areas were calculated in the somas, nuclei, and cytoplasm (regions outside of nuclei but within the soma).

### Whole-genome CRISPR screen next-generation sequencing analysis

21nt long sequencing reads were exported from bcl files using bcl2fastq v2.2.0. These reads were counted by converting them to k-mers and mapping them to a set of 91,536 valid CRISPR library 20nt gRNA sequences. Reads without an exact match in the library were discarded. After merging the sample counts into a count table, the samples were inspected for proper gRNA infection and coverage. With all the samples passing QC, the MAGeCK RRA (Li et al., 2014) was used to perform gene essentiality and enrichment inference. To run MAGeCK, we used the paired samples option, which contained the sorted samples: low GFP sample in control and high GFP samples in the treatment and a set of non-targeting gRNAs as controls. Functional and network analyses of top RBM3 regulator candidates with FDR<0.05 were performed with Metascape (Zhou et al., 2019) and STRING (Szklarczyk et al., 2020).

### RNA-Seq analysis

Fastq files of sequencing reads were processed directly with the nf-core (Ewels et al., 2020) rnaseq pipeline (v3.3) (Patel et al., 2021), mapped to hg38 genome with the gencode v38 annotation (GRCh38.p13). A gene expression quantification table was generated from the salmon output gene counts using the star_salmon aligner option within the nf-core/rnaseq pipeline. Differential expression analysis was performed using DESeq2 (v3.15). Grouped sashimi plots to visualise alternative splicing of RBM3 mRNA between control and cooled i-neurons were generated using rmats2sashimiplot (v2.0.4) and BAM files.

### ENCODE Project data analysis

RBM3 TPM values in RNA-Seq experiments following targeted CRISPR editing in HepG2 and K562 cell lines, together with the matching control samples, were extracted from the publicly available HNRNPH1 (ENCFF039DFP, ENCFF713MXN, ENCFF616BYI, ENCFF586TGE, ENCFF293ODK, ENCFF266YWO, ENCFF053QJC, ENCFF200BWY), SNRNP70 (ENCFF367WUN, ENCFF058OGQ, ENCFF873KLR, ENCFF311ACC), PUF60 (ENCFF682SFK, ENCFF461EMF, ENCFF585BAL, ENCFF643OYR) and KPNB1 (ENCFF819AUU, ENCFF628VDB, ENCFF565TSG, ENCFF073IFA)-knockdown datasets of the ENCODE Project.

To perform splicing analysis on RBM3 mRNA in control and HNRNPH1 knocked-down HepG2 and K562 cells. Raw Fastq files were downloaded from the European Nucleotide Archive (ENA). Fastq files were processed through FastQC (v0.11.8) for quality control. The data quality of each Fastq file was reviewed manually. Reads were mapped to GRCh38.p13 human reference genome by STAR (v2.6.1b). Alignment results were sorted by the coordinate. Output BAM files were indexed by SAMtools (v1.9).

Splicing analysis was performed using rMATS (v4.1.2). Control and knockdown samples were grouped separately to serve as inputs of rMATS. The gene annotation file (GENECODE Human Release 34, GRCh38.p13) was used for analysis. Inclusion levels and FDR values of RBM3 Exon 3a relative to Exon 3 and 4 in the skipped exon output tables (SE.MATS.JCEC) were used to generate bar graphs indicating RBM3 Exon 3a inclusion levels (see Table S3 for corresponding entries). Grouped sashimi plots to visualise alternative splicing of RBM3 Exon 3 and 4 between control and HNRNPH1 knocked down samples were generated using rmats2sashimiplot (v2.0.4) and BAM files.

### HNRNPH1 public iCLIP analysis

Sequence fastq files from public HNRNPH1 iCLIP datasets from HEK293T cells were accessed from ArrayExpress with accession numbers ERR2201859 & ERR2201860 (Braun et al., 2018). Samples were processed with the nf-core/clipseq pipeline v1.0.0 (Ewels et al., 2020), mapped to hg38 genome with the gencode v38 annotation to obtain crosslink counts. Crosslink signal was visualised with library size normalisation and rollmean smoothing (window 5) with clipplotr (Chakrabarti et al., 2021) between exons 3 and 4 of RBM3 (chrX:48575560-48576419:+). RNA G4 prediction was performed with QGRS Mapper using default settings (Kikin et al., 2006).

